# Rapid urban evolution of the dengue mosquito in West African cities

**DOI:** 10.64898/2026.06.25.734560

**Authors:** Michael Amoa-Bosompem, James E Fifer, Dvorah Nelson, Jeffrey K Boateng, Derrick Sackitey, Stephen Oware, Wendegoudi M Ouedraogo, Jewelna Akorli, Athanase Badolo, Noah H Rose

**Affiliations:** Department of Ecology, Behavior, and Evolution, University of California San Diego, La Jolla, CA, USA; Department of Parasitology, Noguchi Memorial Institute for Medical Research, University of Ghana, Accra, Ghana; Laboratoire d’Entomologie Fondamentale et Appliquée, Université Joseph Ki-Zerbo, Ouagadougou, Burkina Faso

**Author notes:** These authors contributed equally to the work (Michael Amoa-Bosompem, James E Fifer).

## Abstract

*Aedes aegypti* is an exceptionally effective global vector of human disease because of its strong specialization on human hosts and habitats. However, in its native range in Africa, many populations never specialized on humans and retained an ancestral generalist ecology. Now, in the rapidly growing cities of Kumasi, Ghana and Ouagadougou, Burkina Faso, we directly document genomic and behavioral evidence of a sudden shift towards greater specialization on humans over just five years. These changes are likely to enhance the ability of these urban mosquito populations to serve as effective vectors of human disease and may have played a role in unprecedented recent outbreaks of dengue fever in West African cities.

## INTRODUCTION

From the earliest days of evolutionary inquiry, a tension has existed around the timescale on which evolutionary processes should be conceptualized (Thompson, 1998). On the one hand, emerging from a broader backdrop of uniformitarian reasoning, an increasing recognition of the deep history of life on Earth gave researchers the framework needed to envision gradual shifts and patterns of diversification over geological time (Mayr, 1982). On the other hand, confronted with the upheavals and Dickensian sprawl of the ongoing industrial revolution, the evolution of a soot-camouflaged morph of the peppered moth became a symbol of the capacity of natural populations to respond remarkably quickly to unprecedented human-driven environmental change (Cook, 2003; Cook et al., 2012). In the ensuing century and a half, it has become increasingly clear that human-driven change is a major evolutionary force – but the consequences of resulting eco-evolutionary change, including its effects on human society, health, and wellbeing, remain incompletely understood (Bay et al., 2017; Johnson & Munshi-South, 2017; Winchell et al., 2023).

The mosquito *Aedes aegypti* is the primary global vector of several important arboviruses, including the viruses that cause Zika (ZIKV), dengue (DENV), chikungunya (CHIKV), and yellow fever (YFV), in large part to its remarkable success in urban habitats across the global tropics (Powell et al., 2018). In these habitats, a tightly coordinated set of traits contribute to the remarkable ability of this species to spread disease (reviewed in (Fifer et al., 2025). These include a strong preference for human host odor (Rose et al., 2023), the ability to make use of artificial container habitats that other mosquitoes might reject (Metz et al., 2023) and a high level of physiological susceptibility to viruses like DENV (Dabo et al., 2024) and ZIKV (Aubry et al., 2020). Perhaps surprisingly given the strength of this specialization, these traits appear to have evolved relatively recently, likely within the last 5,000 years (Rose et al., 2023), with the drying of the Sahara desert and the emergence of a human-specialist niche in human water storage vessels in the hot, dry Sahel region. In fact, in its native African range *Ae. Aegypti* comprises two forms – a widespread ancestral generalist form that is less specialized on human hosts and found across rural and forest habitats, and the human-specialist form which is associated with higher human density areas (Fifer et al., 2026; Rose et al., 2020) and gave rise to global invasives (Crawford et al., 2025; Rose et al., 2023).

In 2018, we found evidence that, beyond the hot, dry habitats of the Sahel, *Ae. aegypti* populations in urban habitats in Africa – particularly the cities of Kumasi, Ghana and Ouagadougou, Burkina Faso, were more attracted to human hosts than nearby rural populations (Rose et al., 2020). These behavioral differences were associated with long tracts of human-specialist ancestry in their genomes, suggesting a recent influx of ancestry in the past few decades (Rose et al., 2020). At the same time, unprecedented outbreaks of dengue fever were documented in these countries and in the city of Ouagadougou (Badolo et al., 2022; Pratt et al., 2026). Taken together, this is suggestive of recent shifts – but urban growth in these cities is still progressing extremely quickly, allowing us to directly test whether urban growth is indeed driving rapid evolutionary change in local mosquito populations. Here we return to these cities five years after our original observations and document a rapid shift towards greater specialization on humans, describe the genomic drivers of this shift, quantify resulting behavioral change, and discuss downstream consequences for the distribution and burden of human disease.

## RESULTS

### Human-specialist ancestry increases with urban growth

First, we sought to test predictions that human specialization in *Ae. aegypti* will increase with human population density (Rose et al., 2020; Fifer et al., 2026). In order to characterize shifts in human specialization associated with urban growth, we sampled *Ae. aegypti* populations in 2023 and 2024 along urban-rural spatial gradients in Burkina Faso (Supp. Fig. 1A) and Ghana (Supp. Fig. 1B) and revisited the growing cities of Ouagadougou and Kumasi sampled in 2018 (Fig. 1). In Burkina Faso, sampling was carried out at two rapidly urbanizing communities in Ouagadougou—Zongo (ZNG) and 1200 Logements (1200L)—as well as the peri-urban locality of Tanghin-Dassouri (TDA) and the rural locality of Kokologo (KLG) (Fig. 1A; Supp. Table 1). In Ghana, we sampled in rapidly urbanizing Kumasi (KUM) -Ahinsan, peri-urban Fumesua (FUM), and rural Bomfobiri (BOM), in addition to 2 communities in Urban Accra – University of Ghana (UG) and Sowutuom (SOW) (Fig. 1B; Supp. Table 1). In total we sampled from nine different communities in 2023/2024, two of which (1200L and KUM) were sampled previously in 2018 (Rose et al., 2020).

**Figure 1.**
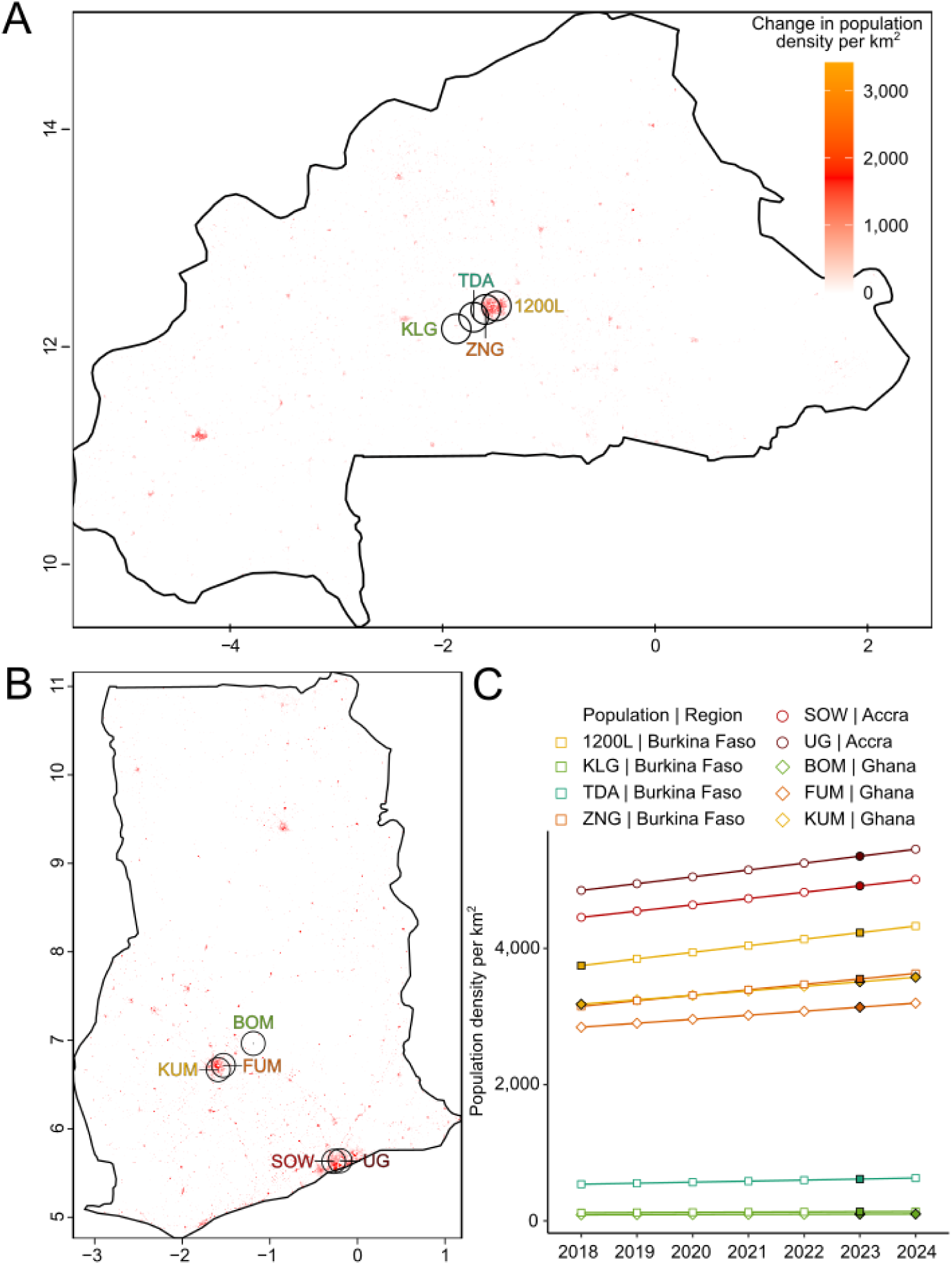
Urban-rural gradients in Ghana and Burkina Faso showing the change in population density per km ^2^ from 2018 to A) 2023 in Burkina Faso and to B) 2024 in Ghana. C) Population density at each year from 2018 to 2024, shapes are colored if *Ae. aegypti* was collected that year. Circles show 15km^2^ buffer area.

We sequenced the whole genomes of 505 individuals collected in the nine communities of Burkina Faso and Ghana and combined this dataset with existing data for 34 genomes at KUM and 1200L from 2018 (Rose et al., 2020). We estimated the proportion of each sample’s genome derived from human-specialist ancestry using FastNGSsadmix to assign structure to the three major genetic ancestry groups present in the native range of *Ae. aegypti*: *Ae. Aegypti formosus* East (*AafE*), *Ae. aegypti formosus* West (*AafW*) and *Ae. aegypti aegypti* (*Aaa i.e.* human-specialist ancestry) (Fifer et al., 2026). We find that genome-wide human-specialist ancestry increased linearly with human population density both in the cities KUM and 1200L from 2018 to 2023 (linear model *p* < 0.001) and along the urban-rural gradients (linear model *p* < 0.001) (Fig. 2A). We also find that human-specialist ancestry increased with urban growth in the regions of the genome most important for predicting human-host preference (hereafter, host-preference outlier regions) (Fig. 2B; linear model temporal *p* < 0.05 and spatial *p* < 0.05).

**Figure 2.**
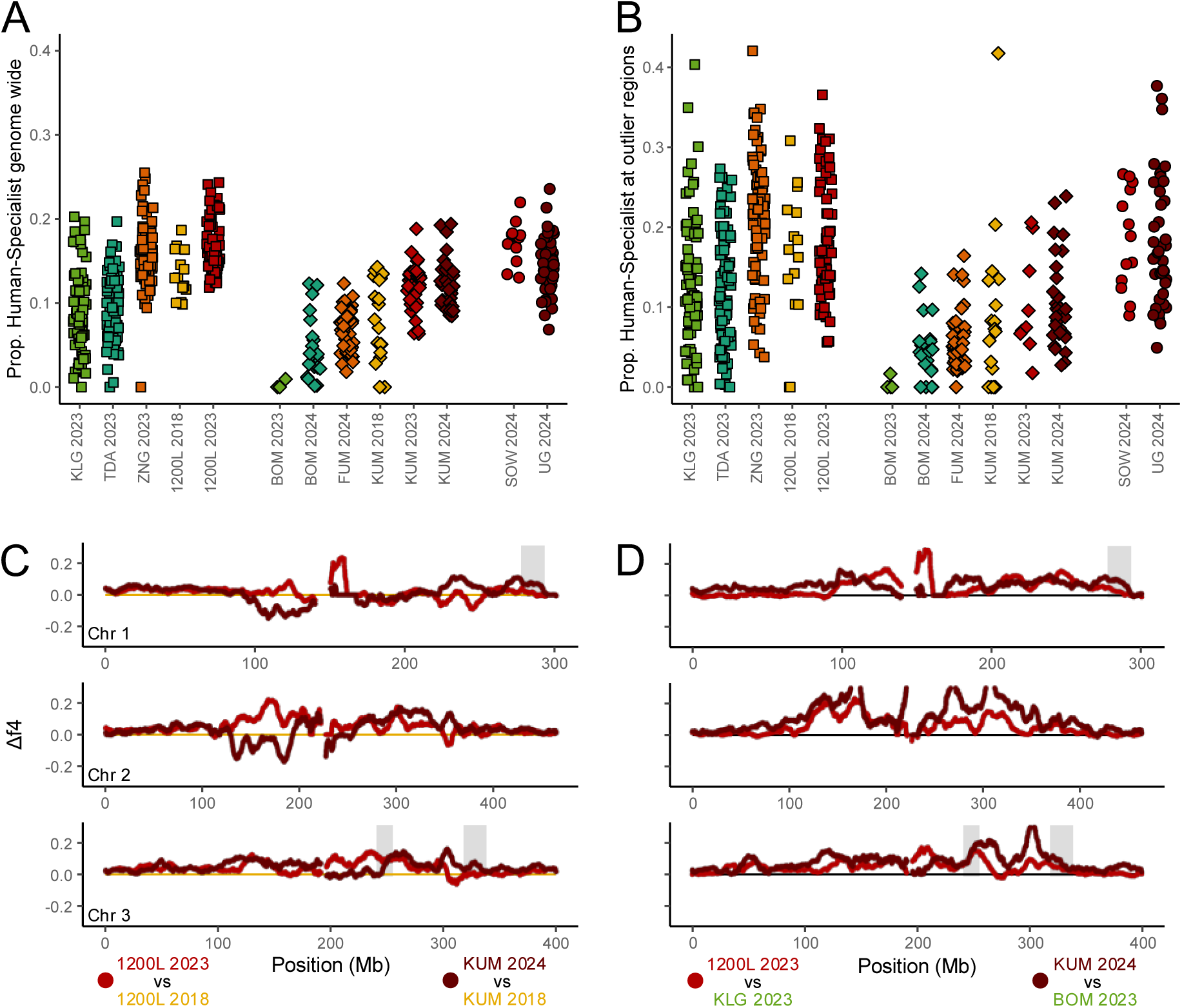
Human-specialist ancestry increases along spatial and temporal axes of urbanization. FastNGSadmix estimates of ancestry A) genome-wide and B) at the host-preference outlier regions from Fifer et al. (2026). C) Temporal and D) spatial estimates of Δf4-ratio calculated in 10mb windows 10kb step. Grey boxes denote host-preference outlier regions from Fifer et al. (2026).

We also used f4-ratio tests to estimate human-specialist ancestry in 10mb windows for all populations (Supp. Fig. 2; Supp. Fig. 3). Both temporal (*i.e.* the change between 2018 and 2023/2024) and spatial (*i.e.* the change in human-specialist ancestry between the most urban and most rural) Δf4 were not significantly enriched at previously identified host-preference outlier regions (Fifer et al., 2026; Supp. Fig. 4), in other words the shifts at the outlier regions do not seem to be significantly greater than the shifts we observe genome-wide. As the host-preference outlier regions were originally defined by calculating the predictive power of ancestry at 10mb windows per a generalized additive model (GAM) and then iterating across R^2^ percentile thresholds and choosing the threshold that maximized the predictive power of all windows passing that threshold per a second GAM, this excludes other genomic regions where human-specialist ancestry still predicts variation in host preference but are non-maximal (in terms of their overall R^2^). To ensure that we are not simply missing enrichment in other genomic windows that are particularly important for host preference, we ran circular permutations for all thresholds tested in Fifer et al. (2026) and found no enrichment in Δf4 for any threshold (Supp. Fig. 5).

### Spatial but not temporal parallelism in shifts between Burkina Faso and Ghana

We investigated parallel signatures in ancestry shifts between four comparisons: Ghana temporal Δf4 versus Burkina Faso temporal Δf4, Ghana spatial Δf4 versus Burkina Faso spatial Δf4, Ghana temporal Δf4 versus Ghana spatial Δf4, and Burkina Faso temporal Δf4 versus Burkina Faso spatial Δf4. We find a signature of parallelization between temporal and spatial Δf4 for Burkina Faso and for spatial Δf4 between Burkina Faso and Ghana, but not for the other comparisons (Supp. Fig. 6; Supp. Fig. 7).

To investigate the divergent temporal signatures between KUM and 1200L, we examined within each location how human-specialist ancestry changed across genomic regions stratified by their initial (2018) frequencies. Specifically, we asked whether temporal shifts were driven by either genomic regions with low initial human-specialist ancestry (∼0) acquiring increased ancestry over time or regions already harboring substantial initial ancestry showing further increases. We find that KUM and 1200L show strikingly different patterns. 1200L exhibited nearly uniform increases in human-specialist ancestry across all genomic windows regardless of initial frequency (slope ≈ 1; Supp. Fig. 8), while KUM showed the largest ancestry increases in regions with initially low frequencies (slope < 1; Supp. Fig. 8).

One intriguing interpretation of these distinct signatures of genomic change over time is that they may reflect two distinct adaptive processes operating at different stages of urban growth. KUM, with lower baseline human-specialist ancestry and population density, appears to be experiencing evolutionary change driven by increasing selection for human-specialist traits as urbanization increases. In this scenario, loci with initially low frequencies (∼0) are furthest from the new equilibrium under strengthened selection and thus experience the largest absolute increases, while loci with higher initial frequencies are already closer to the new equilibrium and show smaller changes. In contrast, 1200L could represent a scenario where strong selection has already been established and ecological dynamics dominate. In this scenario, urbanization might drive increased connectivity to other human-specialist source populations, adding ancestry uniformly across all loci regardless of initial frequency.

To test these interpretations, we modeled both scenarios from first principles using migration-selection balance: p* = 1 -m/s, where p is human-specialist allele frequency, m is gene flow from forest populations, and s is the selection coefficient. For the evolution scenario (KUM), increasing selection (s → 1.2s) yields p* = 1 -m/(1.2s), producing slope < 1 due to boundary effects where under the shifting equilibrium, loci near zero can undergo larger absolute increases than loci that already begin at higher frequencies (Supp. Fig. 9). For the ecology scenario (1200L), increased connectivity adds a uniform pulse of human-specialist ancestry (p* + 0.1), yielding slope ≈ 1 (Supp. Fig. 9). These theoretical predictions match our empirical observations, consistent with the idea that urbanization could result in different genomic signatures in adapting populations, depending on the context and stage of urban growth.

### Urbanization drives shift towards preference for human hosts

We first used a combined host-preference behavioral dataset derived from trials which included newly generated behavioral data from Burkina Faso and Ghana and existing preference data across Africa (Rose et al., 2020; Rose et al., 2023; Fifer et al., 2026) in conjunction with the ecological model from Fifer et al. (2026) to test the optimal human population density buffer size and found a model with a 15km^2^ buffer (as approximately linear) best explained the variation in host preference across Africa (R^2^ = 0.85; Supp. Fig. 10).

Using the genomic-behavioral model from Fifer et al. (2026), we obtained predicted host preference for all populations from human-specialist ancestry at the host-preference outlier regions, which maintained a high correlation (R^2^ = 0.82) with observed preference across both datasets (Supp. Fig. 11). Additionally, predicted host preference does a reasonable job of predicting the observed field host preference index (R^2^ = 0.725; Supp. Fig. 12). We also show that host preference predictions are robust to the method employed to calculate human-specialist ancestry at the outlier regions (*i.e.* f4-ratio traces or fastNGSadmix; Supp. Fig. 13). Within Burkina Faso and Ghana, we found a simple beta regression model with geographic region (*i.e.* Burkina Faso, Ghana or Accra) and a 15km^2^ human population density buffer explains 87.5% of the variation in predicted host preference (Supp. Fig. 14), with human population density as a significant predictor, acting in accordance with the model predictions from Fifer et al. 2026 (Fig. 3B). Further corroborating our findings and suggesting the shift in predicted preference is representative of actual behavioral change, we find observed host preference differs significantly between the sampled years (*p* < 1 × 10^-8^) and between the most urban (KUM, 1200L) and rural (BOM, TDA, KLG) cities in 2023/24 (*p* < 0.01).

**Figure 3.**
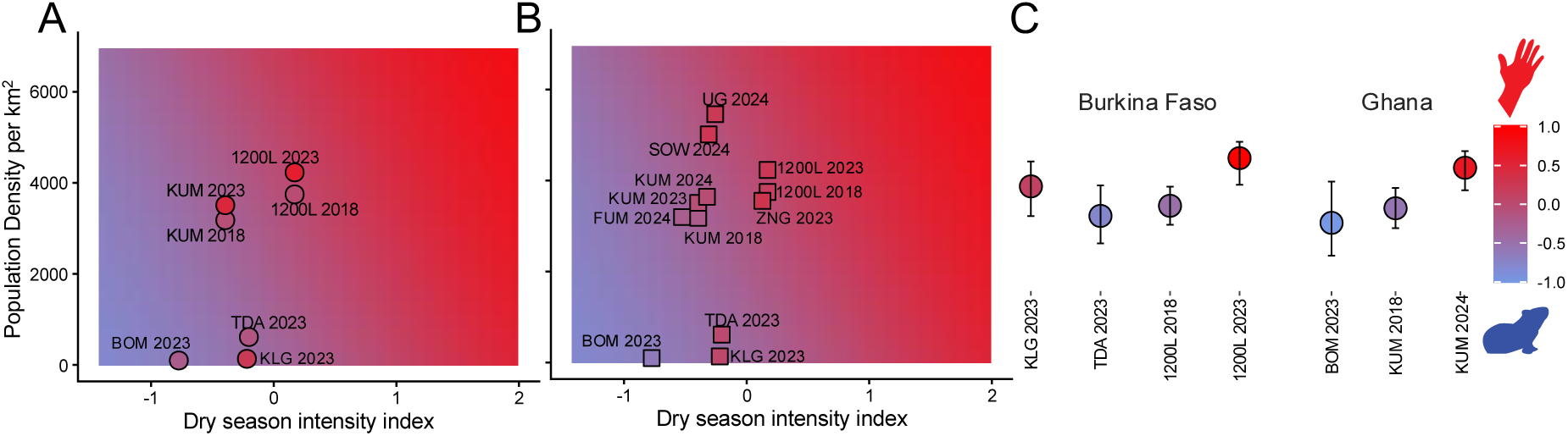
Urbanization results in a shift toward populations with stronger preference for human hosts in Ghana and Burkina Faso. Predicted host preference from GAM in two-dimensional grid space, projected for variations in dry season intensity and population density (15km^2^ buffer) with A) circles representing observed host preference and B) squares showing predicted host preference from the genomic data. C) Observed preference indices organized by country. Vertical line shows 95% confidence intervals.

## DISCUSSION

In this study, we provide evidence supporting previous predictions (Rose et al., 2020; Fifer et al., 2026) that rapid urbanization is driving a shift towards human specialization in cities in sub-Saharan Africa. We find higher levels of human-specialist ancestry in locations with higher human population density. Human-specialist ancestry is linked to a suite of traits putatively associated with *Ae. aegypti* exploitation of environmental niches created by humans (reviewed in Fifer et al., 2025). We further examined one such trait, preference for human host odor, and found direct evidence suggesting that ongoing urbanization is driving greater preference for humans in urban West African populations of *Ae. aegypti*. While all three of our approaches to capture host preference (*i.e.* predicted, observed *ex situ* and observed *in situ*) all broadly align, these are ultimately less direct measures of host feeding behavior compared to blood meal analysis. While comprehensive field studies across urban-rural gradients examining host blood meals are limited and present an important avenue for future work, blood meal analysis in Ouagadougou from 2018 (Ouédraogo et al., 2024) as well as recent Dengue outbreaks in Burkina Faso (Belem et al., 2026) and the first described major dengue outbreak in Ghana in 2024 (Pratt et al., 2026) are all consistent with changing disease transmission dynamics resulting from increased anthropophily in local *Ae. aegypti* populations.

The link between urbanization and human-specialist ancestry is not as straightforward as it may initially appear. The ancestral form of *Ae. aegypti* is an ecological generalist, breeding in a range of natural containers like tree holes and rock pools, and feeding opportunistically on a wide range of non-human animals (Chepkorir et al., 2018; McClelland & Weitz, 1963). This ancestral form likely took advantage of natural habitats that the green Sahara provided prior to the end of the African humid period 5,000 years, a timepoint that coincides with the evolution of the human-specialist form of *Ae. aegypti* (Rose et al., 2023). This major environmental change may have forced *Ae. aegypti* to rely on human stored water to complete its life cycle, kicking off an evolutionary trajectory towards increased reliance on a human-specialist niche (Peterson, 1977; Powell et al., 2018; Rose et al., 2023). Our findings that contemporary urbanization is driving an increase in human-specialist ancestry thus supports a broader pattern emerging from studies of urban evolution (Fifer et al., 2025; Winchell et al., 2023), that the variation which urban selective pressure acts upon can predate modern cities.

The increase in human-specialist ancestry has coincided with an increase in the incidence of *Ae. aegypti*-borne arboviruses such as ZIKV and DENV (Badolo et al., 2022; Pratt et al., 2026). This increase may be attributable to at least two key human specialist-related factors driving vectoral capacity. First, an increase in human-specialist ancestry increases attraction to human hosts, increasing human-mosquito contact (Caldwell et al., 2024; Rose et al., 2020). Other aspects of increased ecological specialization on human habitats (*e.g.* the ability to breed in artificial containers) likely further enhance the effects of human-specialist ancestry on human-mosquito contact (Fifer et al., 2025). Second, human-specialist populations of *Ae. aegypti* show a higher susceptibility to key arboviruses, including ZIKV (Aubry et al., 2020; Caldwell et al., 2024) and DENV, although the DENV effects are complex and appear to vary according to mosquito population and DENV strain (Dabo et al., 2024). This suggests that the progressive increase in human-specialist ancestry in the Sahel requires purposeful and targeted public health mitigation efforts to reduce the potential impact and protect quality of life.

A major focus on urban ecology and evolution studies has been on presenting urban areas as replicated natural experiments to test for parallel evolution (Winchell et al., 2023). The parallel shifts we observe along urban–rural gradients in Burkina Faso and Ghana may be driven less by wholly independent evolution from genetic variation found in nearby forest or rural populations than by connectivity among urban environments. Ongoing gene flow from surrounding forest, rural or other urban populations creates interesting gene-flow selection dynamics in our system. First, ancestry changes are parallel along the urban–rural gradients, with these evolutionary dynamics playing out over a longer timescale that goes back to when the cities were first established (*e.g.* Ouagadougou started experiencing population growth in the early 1900s (Fournet et al., 2008)). In Ouagadougou, five-year temporal changes in ancestry mirror observed spatial clines in ancestry and suggest a regime in which selection is no longer intensifying, and instead expanding urban niche space appears to drive a continued influx of human-specialist ancestry from nearby sources. Kumasi shows a contrasting temporal trajectory, with a signature consistent with increasing selection from 2018 to 2024. Together, these results highlight an alternative explanation for non-parallelism among cities beyond those typically discussed in the literature (Szulkin et al., 2020): even if selective pressures are broadly similar across urban environments, populations may exhibit different adaptive responses because they are at different stages along a shared evolutionary trajectory.

Our results raise the question of how similar dynamics are playing out in other growing cities in Africa. Outside of Ghana and Burkina Faso, other African countries are also dealing with a rising burden of *Ae. aegypti*-borne arboviral disease (Kraemer et al., 2017; Lutomiah et al., 2016; Mordecai et al., 2020; Rose et al., 2022). The role of vector evolution in the emergence of new transmission dynamics is an important topic for future study across the range of this and other important vector species. While we ascribe the lack of temporal parallelism between Kumasi and Ouagadougou to divergence in their stage of urban growth, it is possible that urbanization as a homogeneous selective pressure is an oversimplification and that *Ae. aegypti* populations are also locally adapting to some discrete environmental differences that cannot be parsed without more examples from other cities. In fact, the original context in which the human-specialist ancestry component evolved is not urban at all, instead corresponding to rural environments where human stored water provided the only available aquatic habitat (Rose et al., 2023). This raises the question of the extent to which contemporary urban environments actually provide the same selective pressures, and whether novel or idiosyncratic features in specific cities also play an important role in urban adaptation.

It is not clear what will happen as cities continue to grow, and whether the linear increases in human-specialist ancestry we have described here will plateau as admixed populations become fully urban-adapted. In the native range there is still a large portion of unexplored phenotypic space in the human specialist direction, as invasive lineages demonstrate higher levels of preference for humans than the urban populations studied here (McBride et al., 2014; Rose et al., 2020). However, swamping effects of gene flow from nearby forest environments appear to maintain lower levels of human-specialist ancestry in urban populations in Africa. An informative example can be found in the gene flow-selection dynamics in Rabai, where fully unadmixed human specialist individuals that re-invaded East Africa could be found in indoor domestic habitats prior to 2014 (McBride et al., 2014; Peterson, 1977; Trpis & Hausermann, 1978), but now the pervasive form (or at least as of 2017) has much less human-specialist ancestry and lower human host preference (McBride et al., 2014; Rose et al., 2020). However, given that Rabai is a peri-urban area and there has not been a thorough investigation of *Ae. aegypti* populations in East Africa, it is unclear whether this truly represents some stable plateau. In the invasive range, *Ae. aegypti* populations are unencumbered by gene-flow from the African forest populations, but because they descend from a bottlenecked founding lineage, they also contain less genetic variation for evolution to act on. Despite this, there is some evidence that invasive *Ae. aegypti* populations differ in ecology and host utilization – in South Texas, for example, bloodmeal analysis revealed a higher utilization of dogs than humans (Abdi et al., 2026).

Overall, our results suggest that rapid urban growth is capable of driving rapid evolutionary change in disease vector populations on directly observable timescales of just a few years. In this case, selection seems to act on pre-existing genetic variation from human-specialist populations that existed in a climate-defined niche well before the onset of modern urbanization. Models that assume a stable vectorial capacity in native or long-present species may underestimate the propensity of disease transmission dynamics to shift as humans drive environmental change across global ecosystems.

## METHODS

### Field Collection and Establishment of Laboratory Colonies

This study was conducted across rapidly urbanizing, peri-urban and rural communities 20 -50 Km apart, in Burkina Faso and Ghana – in 2023 and 2023/2024, respectively. We collected various combinations of mosquito eggs and adults from each community (Supp. Table 1). Mosquito eggs were collected with ovitraps made of 32 oz black plastic cups (Karat Packaging, Chino, CA), with holes in the side, lined with seed germination paper (Anchor, St. Paul, MN), and filled with water to cover half the germination paper. Ovitraps were placed in/by tree holes, artificial containers, ponds, nurseries/gardens, occupied buildings, uncompleted buildings and at the edge of forests for 2 -7 days. Germination paper with eggs were allowed to air dry and stored in airtight whirl-pak bags for transportation.

Mosquito eggs on germination paper were hatched in separate pans of hatch broth. Hatch broth was made by dissolving a quarter ground TetraMin Tropical Tablets fish food (Tetra, Blacksburg, VA) in 1 liter of deoxygenated water. Eggs once hatched were reared to adults. Each colony was established from mosquitoes collected from one ovitrap, with at least 2 colonies for each sampling site. Established colonies were used in host odor behavior testing downstream.

Adult mosquitoes were collected with paired human-sheep-host-baited BG pro or BG sentinel mosquito traps (Biogents, Cary, NC) paired with yeast for the Ghana collections. Black round-neck T-shirts slept in overnight were used for human bait while sheep hair stored in -20 °C and transported in airtight Ziplock bags was used as non-human bait. Human and sheep baited traps were placed about 20 cm apart. Traps were set overnight from late afternoon. At least 2 traps were set at each collection site for 4-7 days. Mosquitoes collected each day were separated by location, trap, human-bait (Supp. File 1), genus and sex, submerged in RNAlater (Fisher Scientific, Waltham MA) and stored at -20 °C until transported with ice packs in insulated backpacks (and stored in -20 °C until processed).

### Host Odor Behavior Testing

Host odor behavior was tested using a previously described method with modifications (Rose et al., 2020). Briefly, 7-to 14-day old mosquitoes (g2 -g13) were starved of sugar overnight to prime them for feeding. A custom-made two-port olfactometer with valves for air circulation and CO2 supply was used. Inflow and outflow were set to a rate of approximately 0.3 m/s. The olfactometer had two host chambers; one held a guinea pig (*Cavia porcellus),* and the other contained the forearm of a volunteer (28 year old, European-American female). Human subjects breathed gently into the chamber through the opening every 30s to match the animal’s breath. In each trial 30-135 female mosquitoes were allowed 5-10 minutes to acclimate before valves were opened for 10 minutes for air circulation. Mosquitoes choosing to fly upwind toward each host odor were trapped and counted. Two female guinea pigs were used on alternating days, and the side of the human versus guinea pig was switched every two days.

Populations (*i.e.* the location origin and year of collection) were only included in downstream analyses if sample size allowed for > 5 trials, after removing trials that had high mortality (> 50%). This resulted in the following populations retained for downstream analyses: 1200L 2023 (g5), 1200L 2018 (g13), KUM 2023 (g5), KUM 2018 (g13), BOM 2023 (g3), KLG 2023 (g4), and TDA 2023 (g4). KUM 2018 and 1200L 2018 were later generations of the same colonies used previously in Rose et al. (2020) (Supp. Fig. 15A) Each colony was separated into 5 trials on day 0 and tested iteratively for 5 days. We also used two control lab strains from previous olfactometer studies (Rose et al., 2020; Fifer et al., 2026), a human specialist ORL (g27) and a generalist U52 (g23), each tested for 6 days (Supp. Fig. 15A). Response rate was between 20-80% (Supp. Fig. 15B), which aligned with expected rates based on previous behavioral trials (Rose et al., 2020).

Host preference was estimated in R (R Core Team, 2024) with the package glmmTMB (Brooks et al., 2017) using a generalized linear mixed model with a beta binomial distribution to account for over dispersion, as previously described (Rose et al., 2020, Fifer et al., 2026). 2020 behavioral data from (Rose et al., 2020) for KUM 2018 and OGD 2018 were also included in the analysis. The collection years and sample origins were included as fixed effects, and the populations and trial day were included as random effects, to account for variation among populations and trial days. Probability of host preference using 95% confidence intervals were estimated using emmeans (Lenth & Piaskowski, 2017) and the confint function from the package mass (Ripley et al., 2013) in R. Probability (p) of choosing a human host was then transformed into a preference index with a range of –1 to 1 (formula: 2p-1), where an index of zero means mosquitoes from that population were equally likely to choose either host. We also collected a “field preference index” for populations BOM, FUM, UG, SOW and KUM in 2024 and TDA, KLG, 1200L and ZNG in 2023 from the paired human and sheep baited BG traps in the same manner as described above for host preference index.

### Whole Genome Resequencing and Variant Calling

We extracted total nucleic acid (TNA) from 24 – 48 bait-captured mosquitoes, from each collection site, depending on yield (Supp. File 1), G0’s from established mosquito colonies, and field collected larvae, using the Bio-On-Magnetic-Beads (BOMB) TNA extraction protocol with modifications (Supp. File 1). Briefly, mosquitoes were individually homogenized in 1X GITC (MilliporeSIGMA, Darmstadt, Germany), treated with Isopropanol (MilliporeSIGMA, Darmstadt, Germany) and captured with carboxyl-coated magnetic beads (Fisher Scientific, Waltham MA) in TE buffer (Fisher Scientific, Waltham MA), before eluting in nuclease free water (Fisher Scientific, Waltham MA). DNA extractions were prepared for low-coverage whole genome sequencing (lcWGS) using a TN5 library protocol (Langdon et al., 2022) and sequenced to an average of 1.2X (Supp. Fig. 16) with PE 151 bp reads using the Illumina Novaseq X plus 150PE platform. We combined this new lcWGS dataset from 2023–2024 with previously generated ∼10–15× whole-genome sequencing data for *Ae. aegypti* collected in 2018 from 1200L and KUM (Rose et al., 2020).

All sequences were mapped to a version of the L5 reference genome (Matthews et al., 2018) that was updated with WGS data from 100 unrelated African male *Ae. aegypti* (Rose et al., 2020). Reads were trimmed using TRIMMOMATIC [ILLUMINACLIP:NexteraPE-PE.fa:2:30:10:2:keepBothReads LEADING:3 TRAILING:3 MINLEN:36] mapped to the reference using bwa (Li & Durbin, 2009) converted to bams and filtered to only include reads with mapping quality >20 with samtools (Li et al., 2009), duplicates removed using picard (“Picard Toolkit,” 2019) and clipping overlaps with BamUtil’s clipOverlap (Jun et al., 2015).

We used ANGSD’s likelihood-based calling methods (Korneliussen et al., 2014) for all genotype calling. We called genotypes with filters -minMapQ 10 -minQ 20 using a previously identified set 14,045,728 high confidence SNP calls (Rose et al., 2020) for major and minor allele classifications. Related individuals were filtered out using ngsRelate (Korneliussen & Moltke, 2015), using the coefficient of kinship estimate to remove putative first cousins (theta = 0.0625). *Ae. aegypti* species identity was confirmed using a PCoA created via a covariance matrix based on single-read resampling calculated in ANGSD (Supp. Fig. 17).

### Estimating human-specialist ancestry

We estimated population level ancestry using both FastNGSadmix (Jørsboe et al., 2017) and f4-ratio tests. We used FastNGSadmix with a 18,434,131 locus panel, previously generated to accommodate a wide range (10x-0.0001x) of genomic coverage (Fifer et al., 2026), to estimate each sample’s assignment to the three major genetic ancestry groups present in *Ae. aegypti*: *Ae. aegypti formosus* East (*AafE*), *Ae. aegypti formosus* West (*AafW*) and *Aaa* (*i.e.* human-specialist ancestry). We also used windowed f4-ratio tests (10mb window, 10kb step) on population level allele frequencies to estimate human-specialist ancestry. For f4-ratio tests we used a tree with unadmixed *AafW*, unadmixed *Aaa*, query population, unadmixed *Aaa*, and unadmixed *AafE* as the outgroup following Fifer et al. (2026). We first analyzed the categories 1) BG with animal bait 2) BG with human bait and 3) oviposition collections, as three distinct subgroups and tested whether there was divergence in ancestry estimates across the three collection methods using a linear model. We find no influence of trap type on variation in human-specialist ancestry per a linear model lm (human-specialist ancestry ∼ collection method) (*p* = 0.272) using the car package (Fox & Weisberg, 2018) in R and so combined all trap types for downstream analyses.

Lastly in addition to whole-genome estimates of ancestry we also calculated ancestry at genomic regions previously shown to be particularly important for predicting host preference (Fifer et al., 2026), hereafter “host-preference outlier regions”, again estimating with both fastNGSadmix and f4-ratio tests.

### Investigating parallel shifts in ancestry with increased urbanization

For testing whether shifts in the proportion of human-specialist ancestry genome-wide or at host-preference outlier regions were associated with urban growth, we used whole-genome or outlier region estimates from fastNGSadmix as we are able to go beyond the population level resolution estimates from f4-ratio ratios and obtain estimates for individuals. We looked for temporal and spatial associations between human-specialist ancestry and urbanization using two different linear models, “human-specialist ancestry ∼ time” to compare 2018 sampling with 2023/2024 and “human-specialist ancestry ∼ time” to compare between the most urban (1200L, KUM) and the most rural (KLG, BOM) communities.

While fastNGSadmix allows for individual level estimates, it is limited to genomically broad estimates. To test how urbanization impacts shifts in ancestry on more genomically local levels we use f4-ratio estimates, calculating Δf4 by finding the difference in raw f4-ratio values between the population of interest and some reference population. For temporal comparisons we calculated Δ f4 as the shift in ancestry between 2018 and 2023 or 2024, for 1200L and KUM respectively. We also calculated spatial Δf4 by comparing the difference in f4-ratio between the most urban (1200L, KUM) and the most rural (KLG, BOM) for 2023 Burkina Faso and 2024 Ghana data. We investigated whether temporal or spatial Δ f4 was enriched in outlier regions with a circular permutation test (5,000 permutations).

We investigated parallelism between and within Burkina Faso and Ghana both in temporal and spatial Δf4 shifts in ancestry with a pearson correlation test, testing the correlation in Δf4 shifts across all 10mb windows (non-overlapping) and assessing significance by comparing observed correlations with a circular permutation null (10,000 permutations).

Within each location we examined whether temporal shifts were driven by either genomic regions with low initial human-specialist ancestry (∼0) acquiring increased ancestry over time or regions already harboring substantial initial ancestry showing further increases. We compared the slope of a regression line plotting non-overlapping windows of ancestry for 2018 versus 2023/24, separately for both 1200L and KUM. To understand these slopes in the context of broader ecological and evolutionary processes, we simulate two different scenarios that we believe to be relevant for our populations. For both scenarios we start with the equation for migration-selection balance: p* = 1 -m/s, where p is human-specialist allele frequency, m is gene flow from forest populations, and s is the selection coefficient. In both cases, we use this as a baseline to represent the 2018 populations and iterate across 2,000 values of *s* (0.001–1), setting *m* = 0.01. We simulate a scenario of increasing selection from urbanization (s → 1.2s) yielding p* = 1 -m/(1.2s) and a second scenario of increased connectivity from other human-specialist population sources, treating increased connectivity as a uniform pulse of human-specialist ancestry (p* + 0.1).

### Host preference shifts associated with urbanization

Human population density was calculated using the 2025a WorldPop 100m constrained estimates (Bondarenko et al., 2025) converting to mean density of ppl/km^2^ for buffer sizes of 5km^2^, 10km^2^, 15km^2^, 20km^2^, 25km^2^, 30km^2^, 35km^2^, 40km^2^, 45km^2^, 50km^2^, 60km^2^, 70km^2^, 80km^2^, 90km^2^ and 100km^2^. We combined observed preference data here with observed preference data from Fifer et al. (2026) to test the optimal buffer size for explaining how host preference was partitioned across the native range of *Ae. aegypti* using the ecological model from Fifer et al. (2026). We did this by comparing generalized additive models (GAMs) via the R package mgcv (Wood, 2017) using a model of preference ∼ buffer + dry season intensity variables + batch, where batch indicated observed preference from the current project or Fifer et al. (2026). The dry season intensity variables consistent of bioclim2 variables mean temperature of driest quarter (bio9), precipitation seasonality (bio15), precipitation of driest quarter (bio17) and precipitation of coldest quarter (bio19), CHELSA variables net primary productivity (npp) and maximum potential evapotranspiration (pet penman max) (Fifer et al., 2026). For each GAM we used a beta regression family with the shrinkage smoother “ts” to allow for non-linear associations between ancestry and our behavioral data. We rescaled preference to be between 0 and 1 ((Preference + 1) / 2) to deal with data frequently approaching -1 and accommodate beta regression assumptions. We selected the buffer based on AIC and maximizing R^2^. We used likelihood ratio tests to compare a model with location (*i.e.* urban *vs* rural) or year (*i.e.* 2018 *vs* 2023, urban only) to a null model accounting for day-to-day and community variation.

Using the genomic-behavioral model from Fifer et al. (2026), we obtained predicted host preference for all populations from human-specialist ancestry at the host-preference outlier regions (windows starting at chromosome 1 position 279,100,000, chromosome 3 position 319,820,000 and chromosome 3 position 326,440,000; Fifer et al. (2026)). We evaluated the correlation between host preference predictions generated using human-specialist ancestry at outlier regions inferred by f4-ratio traces versus fastNGSadmix, to test whether predictions are robust to ancestry inference methodology. To test whether urbanization is associated with changes in host preference in Burkina Faso and Ghana, we used the beta regression model “predicted host preference” ∼ human population density + region, where region refers to either Burkina Faso or Ghana. We selected the population density buffer here from the optimal buffer determined from the full observed preference GAM described above.

To derive a univariate “Dry Season Intensity”, we extracted the term-wise partial contributions of the six climatic smooths from the fitted GAM on the model link scale and summed them for each observation. This summed quantity represents the combined contribution of dry season intensity to host preference, independent of the population density term. To visualize the joint effects of dry season intensity and population density in reduced dimensions, we fit a second GAM using the linear predictor from the full model as the response and modeled it as a function of the derived climatic index, population density, and batch. This reduced model provided a smooth two-dimensional approximation to the original fitted surface while preserving differences between batches. For visualization, predictions were generated for the Fifer et al. (2026) batch specifically and then transformed back to the response scale using the inverse link function from the original beta-regression model.

## ACKNOWLEDGEMENTS

This work was initiated under NIH grant K22AI166268 to NHR and completed with support from NIH R35GM159987 to NHR. JEF was supported by a Helen Hay Whitney Foundation Postdoctoral Fellowship. NHR received support from a Hellman Fellowship, and DN received support from a Curci Fellowship.

## SUPPLEMENTARY MATERIAL

**Supplemental Table 1.**
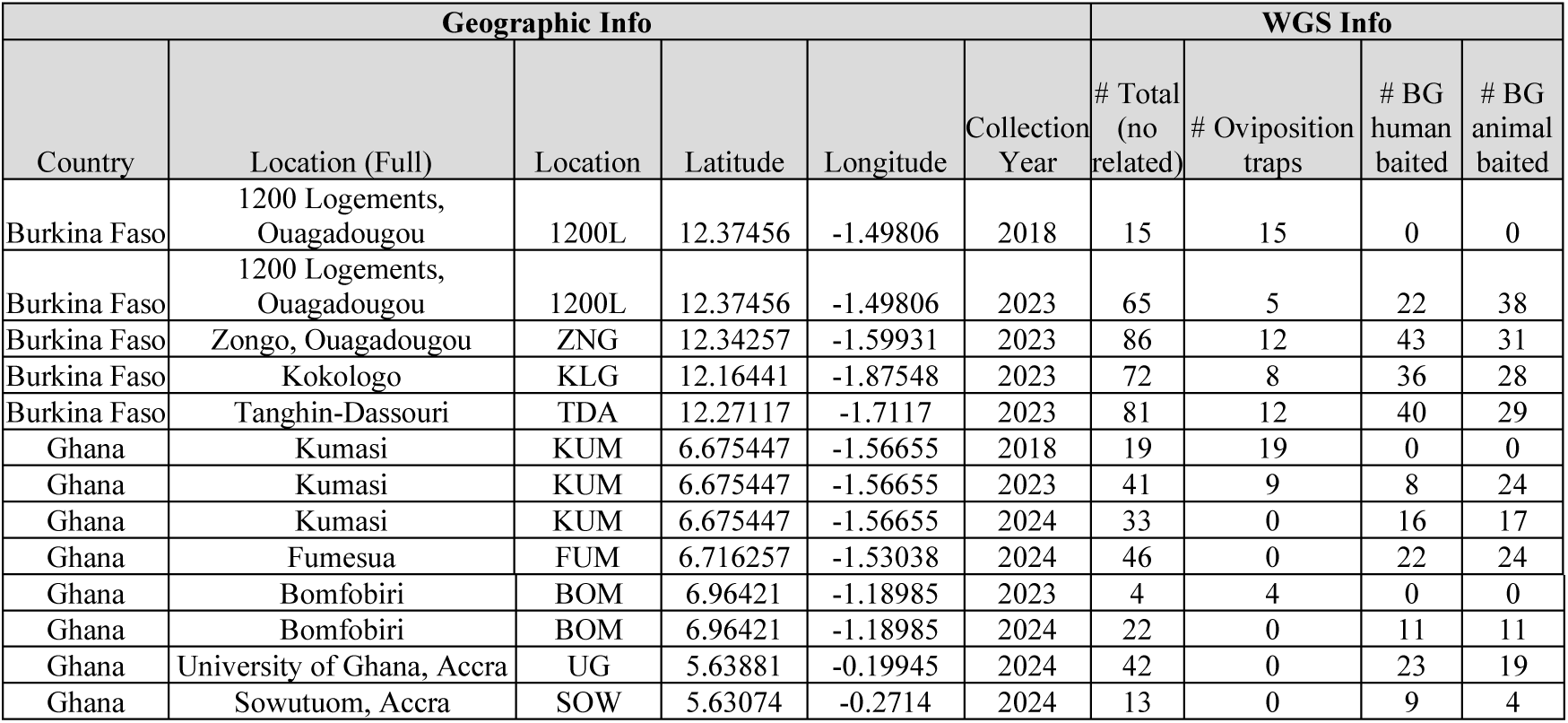
Burina Faso and Ghana populations metadata. 1200L 2018 and KUM 2018 are from Rose et al. (2020).

**Supplemental Figure 1.**
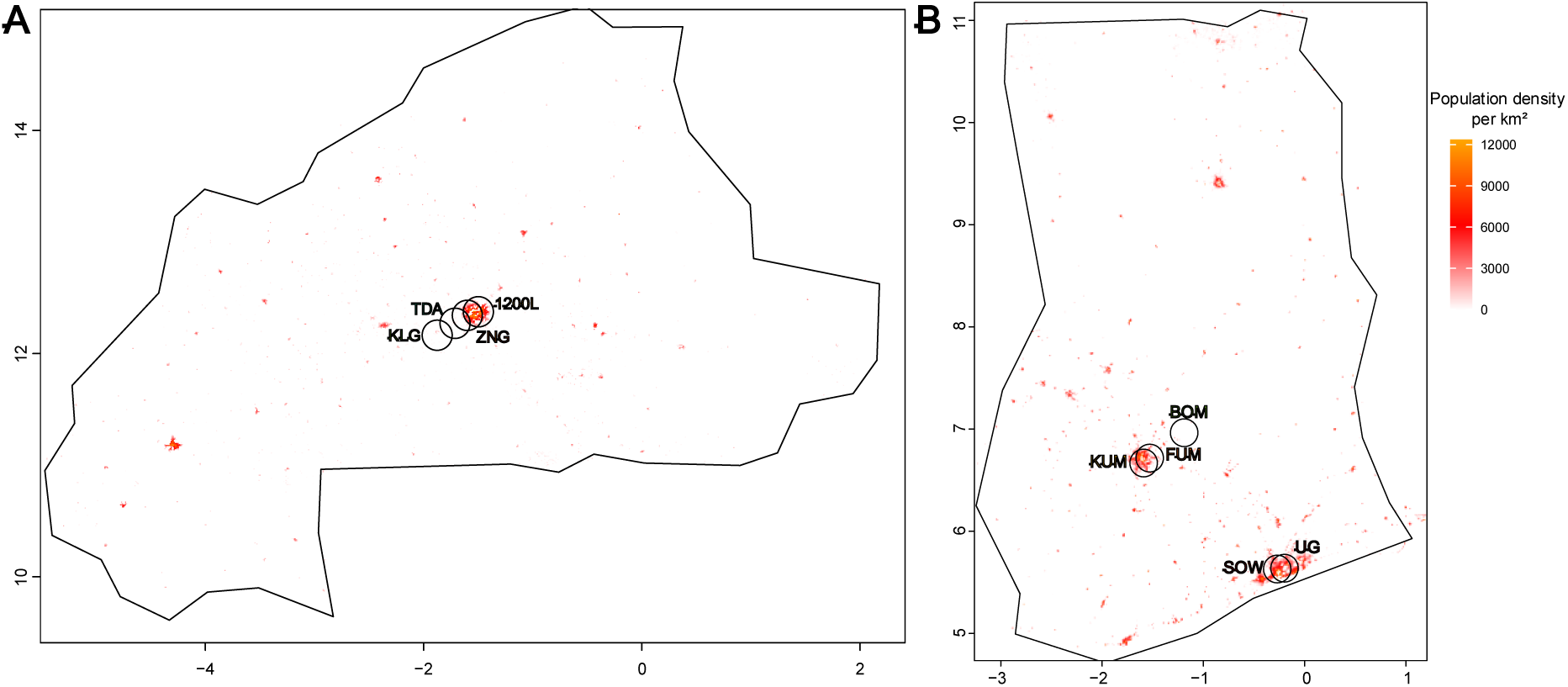
Contemporary (from most recent year sampled – 2023 or 2024) population density per km^2^ for A) Burkina Faso and B) Ghana. Circles show 10km^2^ buffer.

**Supplemental Figure 2.**
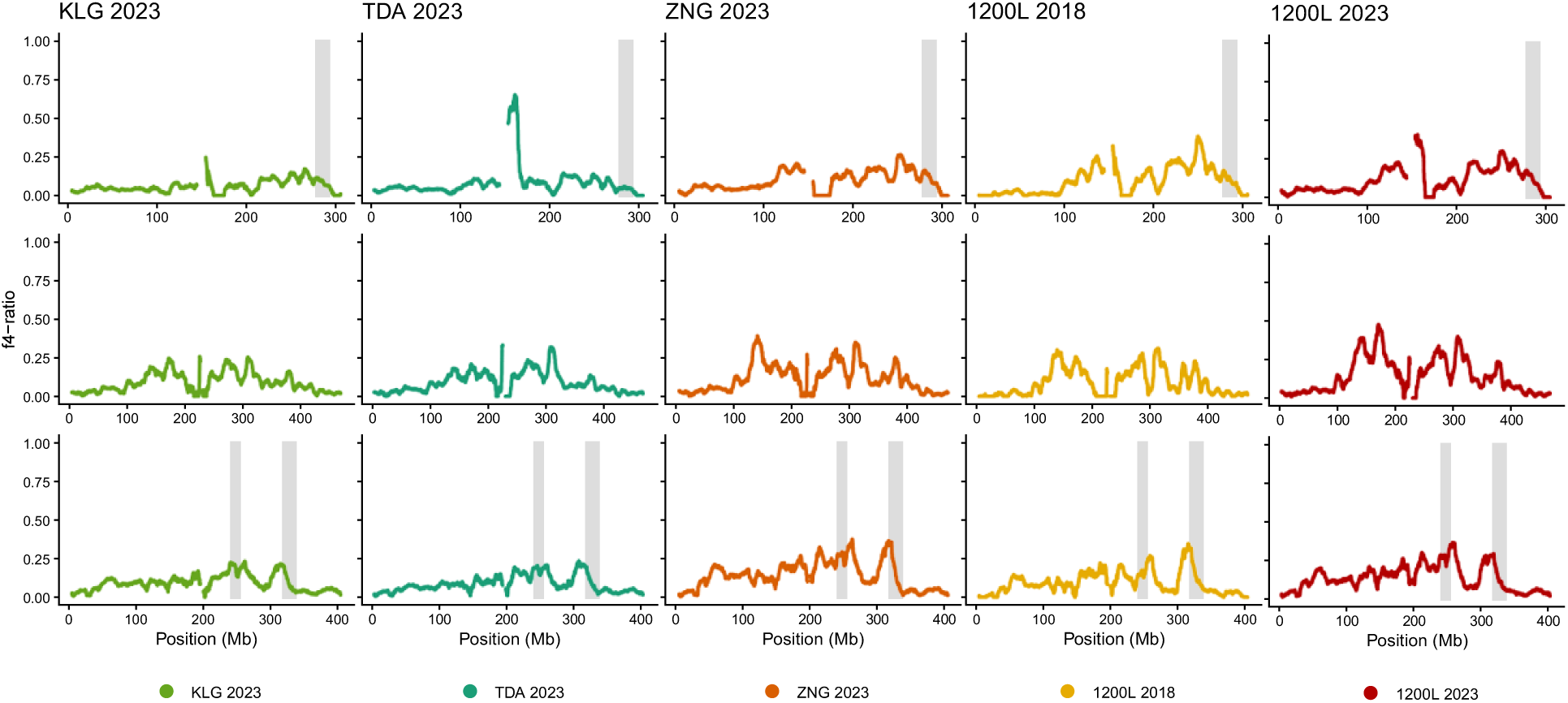
Burkina Faso human-specialist ancestry trace calculated in 10mb windows using f4-ratio estimates. Grey boxes: host-preference outlier regions. Top to bottom: chromosomes 1,2,3.

**Supplemental Figure 3.**
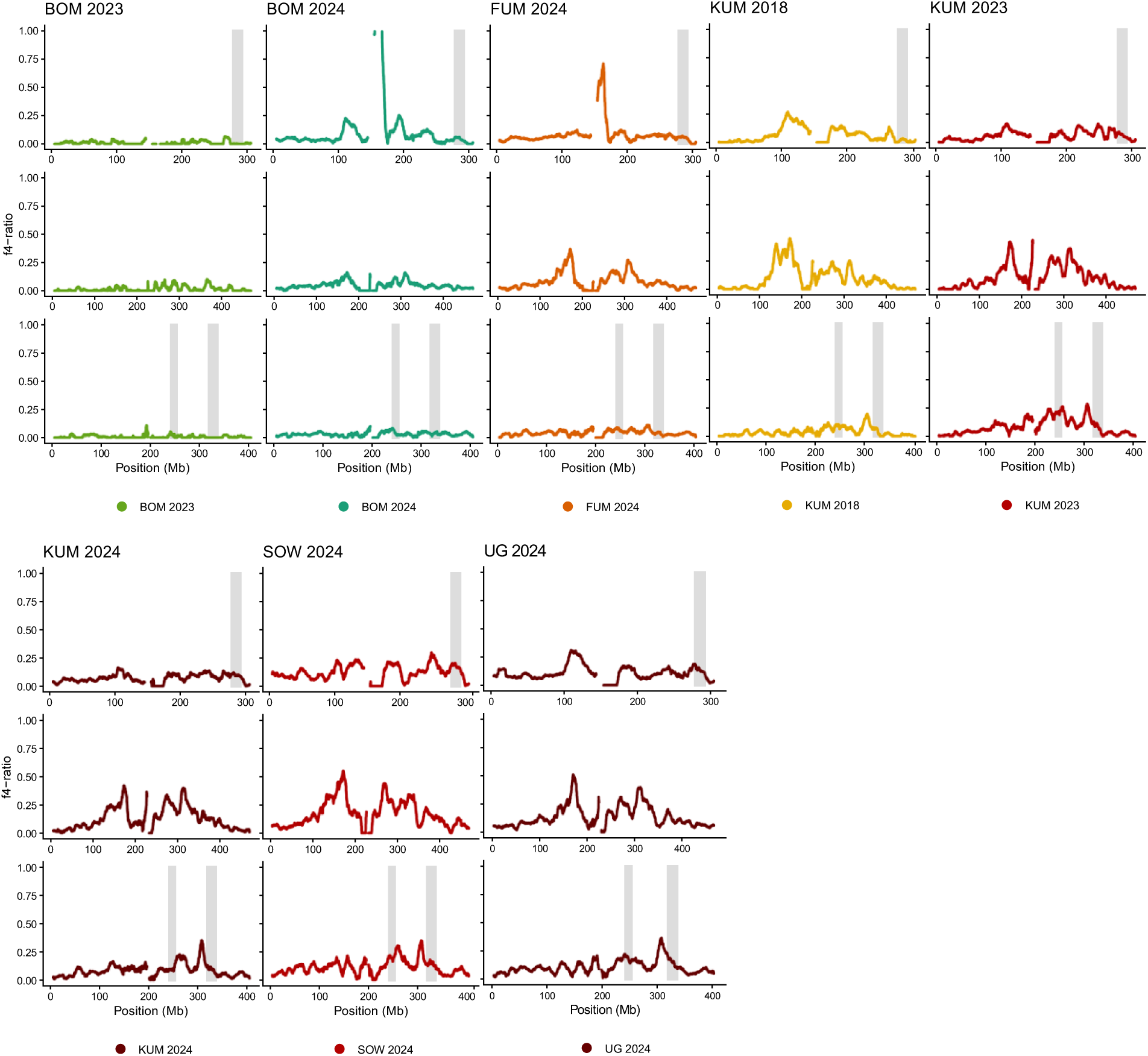
Ghana human-specialist ancestry trace calculated in 10mb windows using f4-ratio estimates. Grey boxes: host-preference outlier regions. Top to bottom: chromosomes 1,2,3.

**Supplemental Figure 4.**
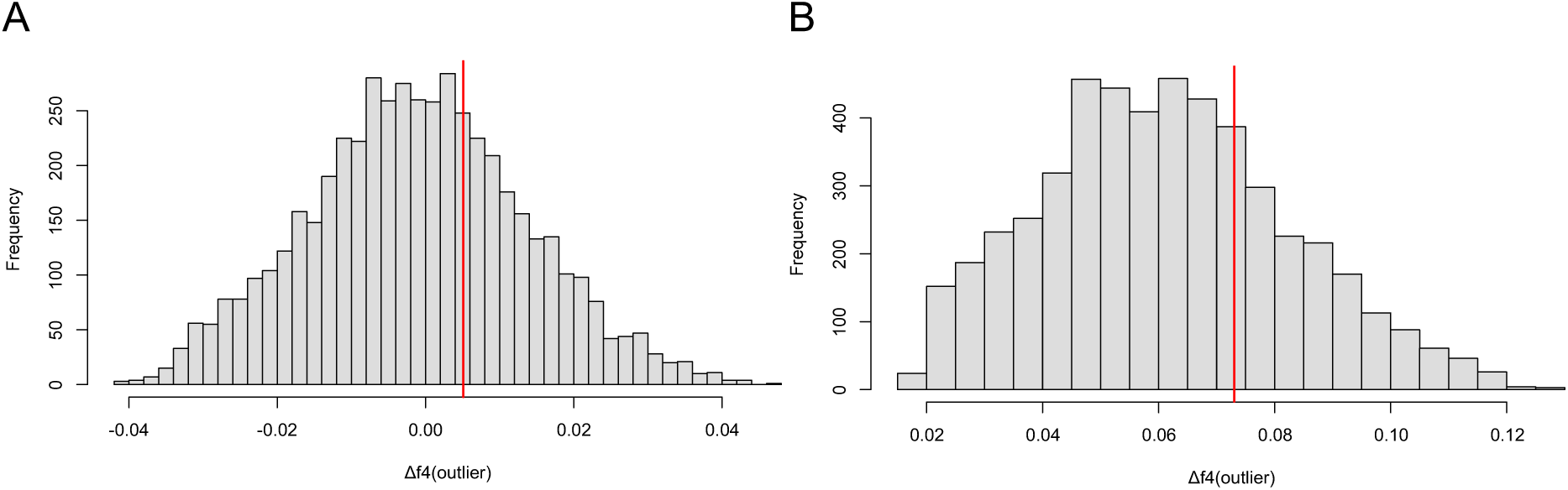
Circulation permutation test shows no enrichment of Δf4-ratio within outlier regions for both A) temporal and B) spatial Δf4-ratio. Vertical red line shows observed Δf4-ratio, grey histogram shows null distribution.

**Supplementary Figure 5.**
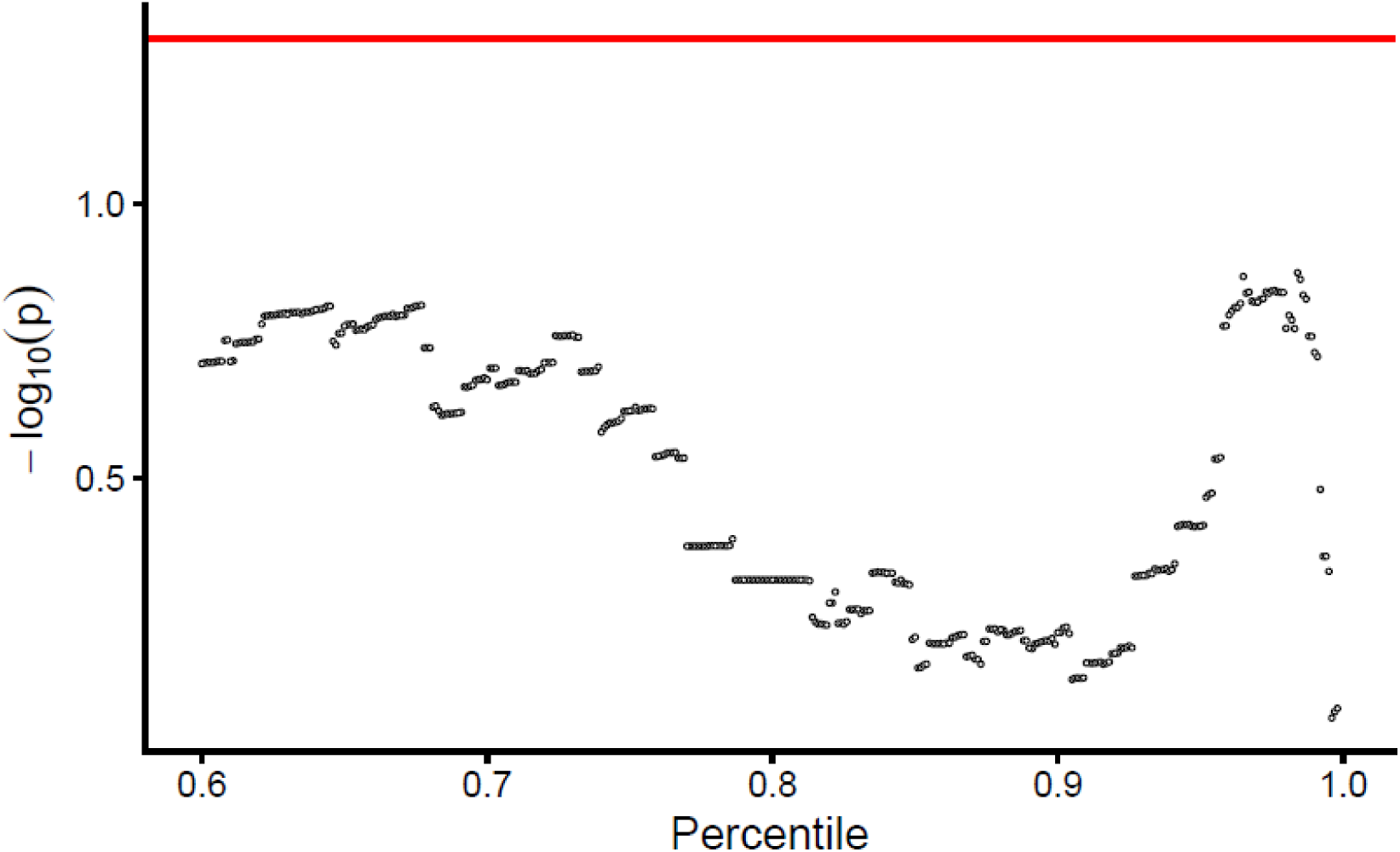
Plot shows p-values from circular permutation testing of delta f4 enrichment of outlier regions, where outlier regions are defined according to different R^2^ percentile thresholds (0.60-0.999, increasing by 0.001) from Gaussian GAMs on logit transformed preference data in Fifer et al. (2026). Horizontal red line denotes upper limit for p-value significance (0.05).

**Supplementary Figure 6.**
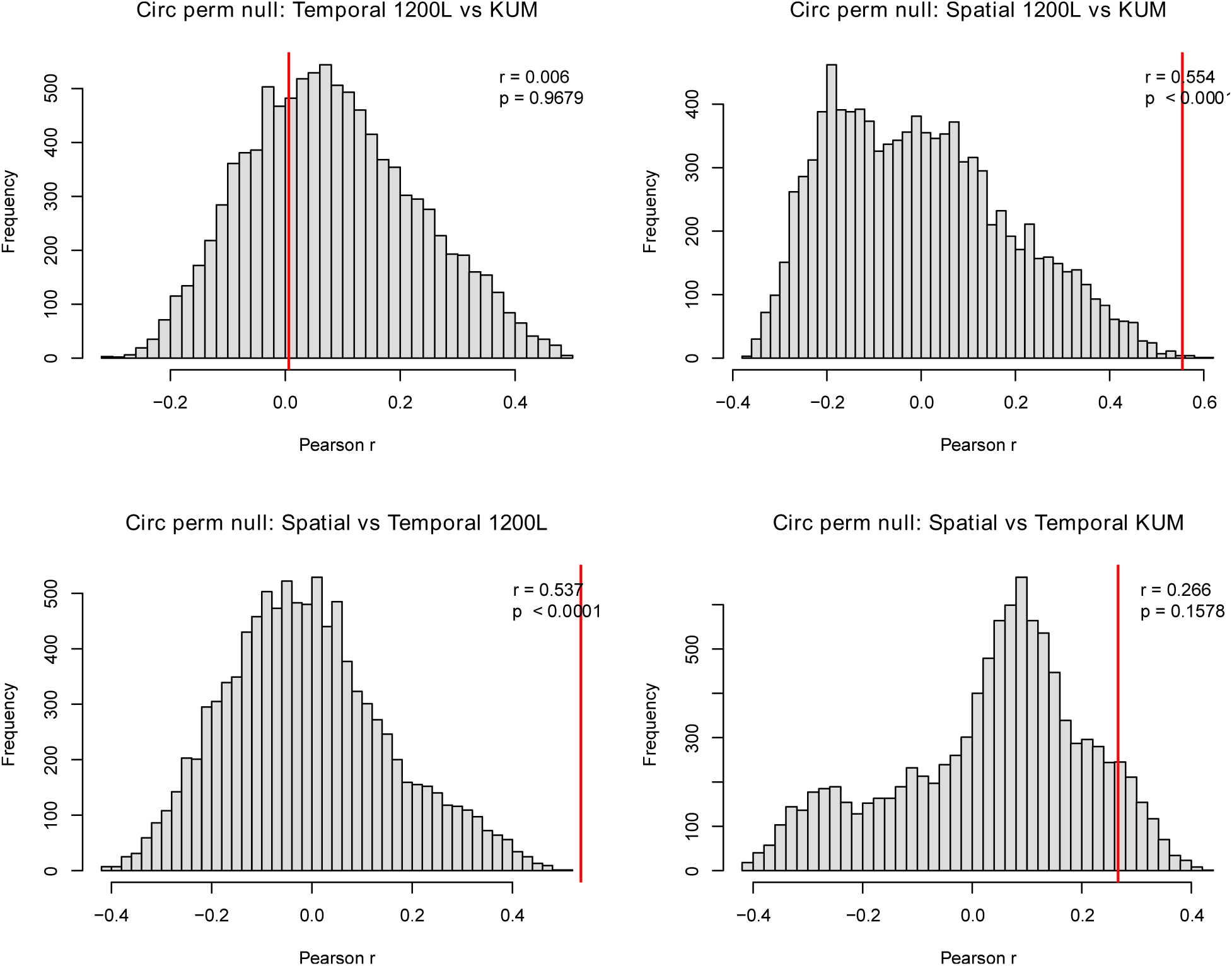
Circulation permutation testing parallel shifts in Δf4-ratio through pearson correlation of all non-overlapping windows. Vertical red line shows observed pearson r, grey histogram shows null distribution generation from 10,000 permutations.

**Supplementary Figure 7.**
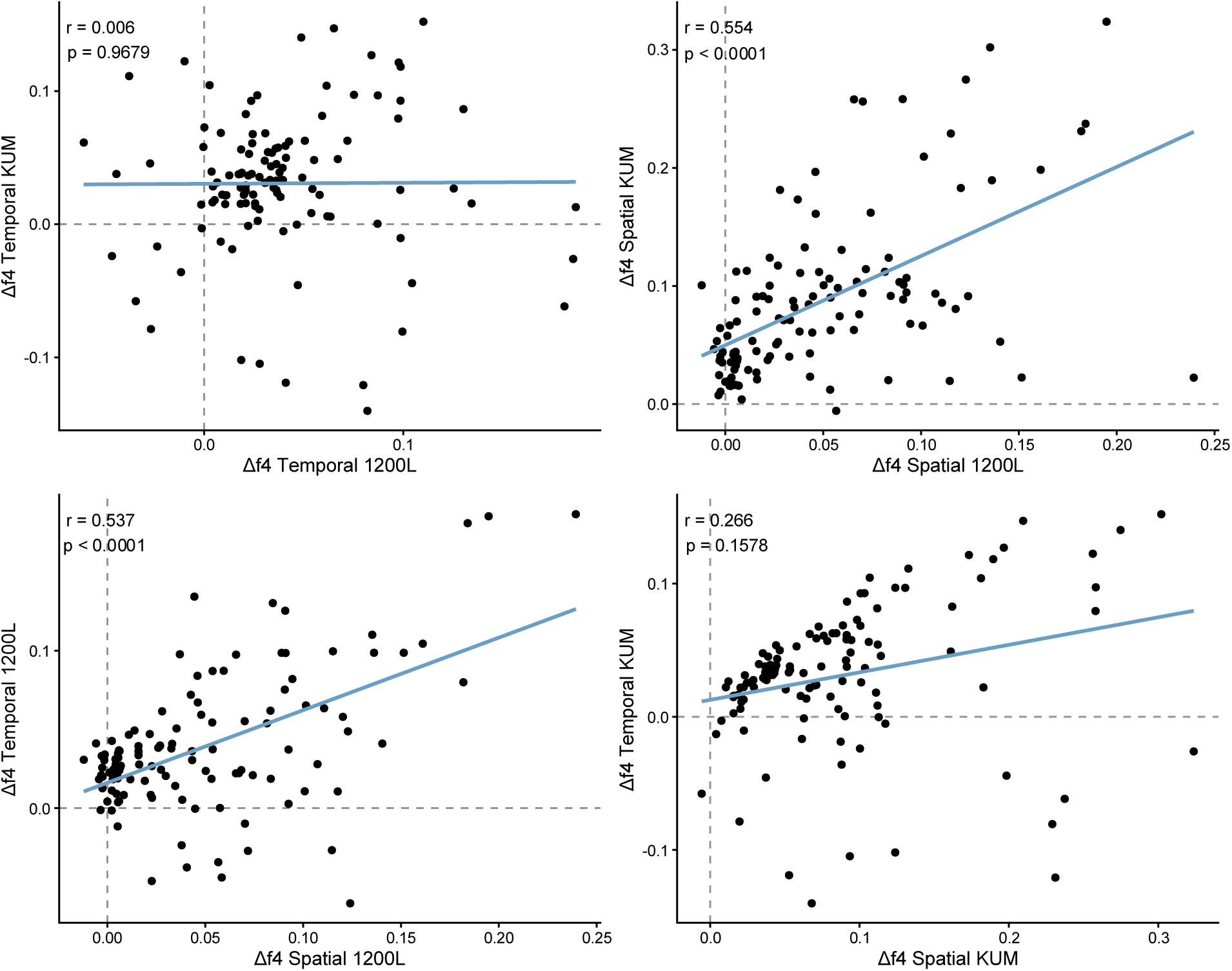
Plot investigating parallel shifts of Δf4 for four separate comparisons. Each dot represents Δf4 at non-overlapping 10mb window. P-values generated through circulation permutation test (Supp. Fig. 6), r: observed pearson’s r. Blue line shows linear regression fit.

**Supplementary Figure 8.**
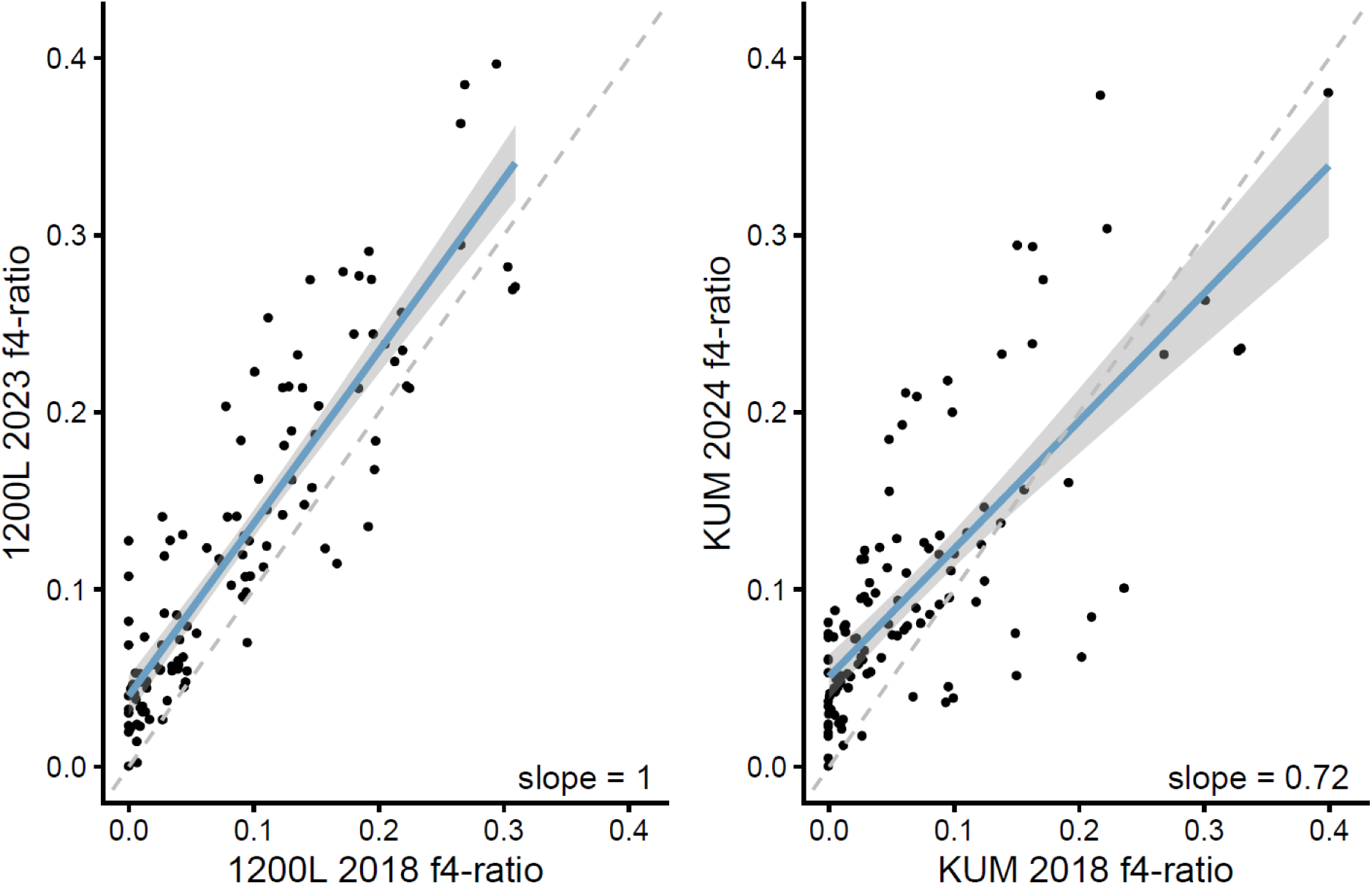
Temporal shifts in human-specialist ancestry reveal contrasting genomic dynamics between Burkina Faso and Ghana. Each dot represents f4-ratio value at non-overlapping 10mb window. Blue line shows linear regression fit with slope rounded to 2 decimal places.

**Supplementary Figure 9.**
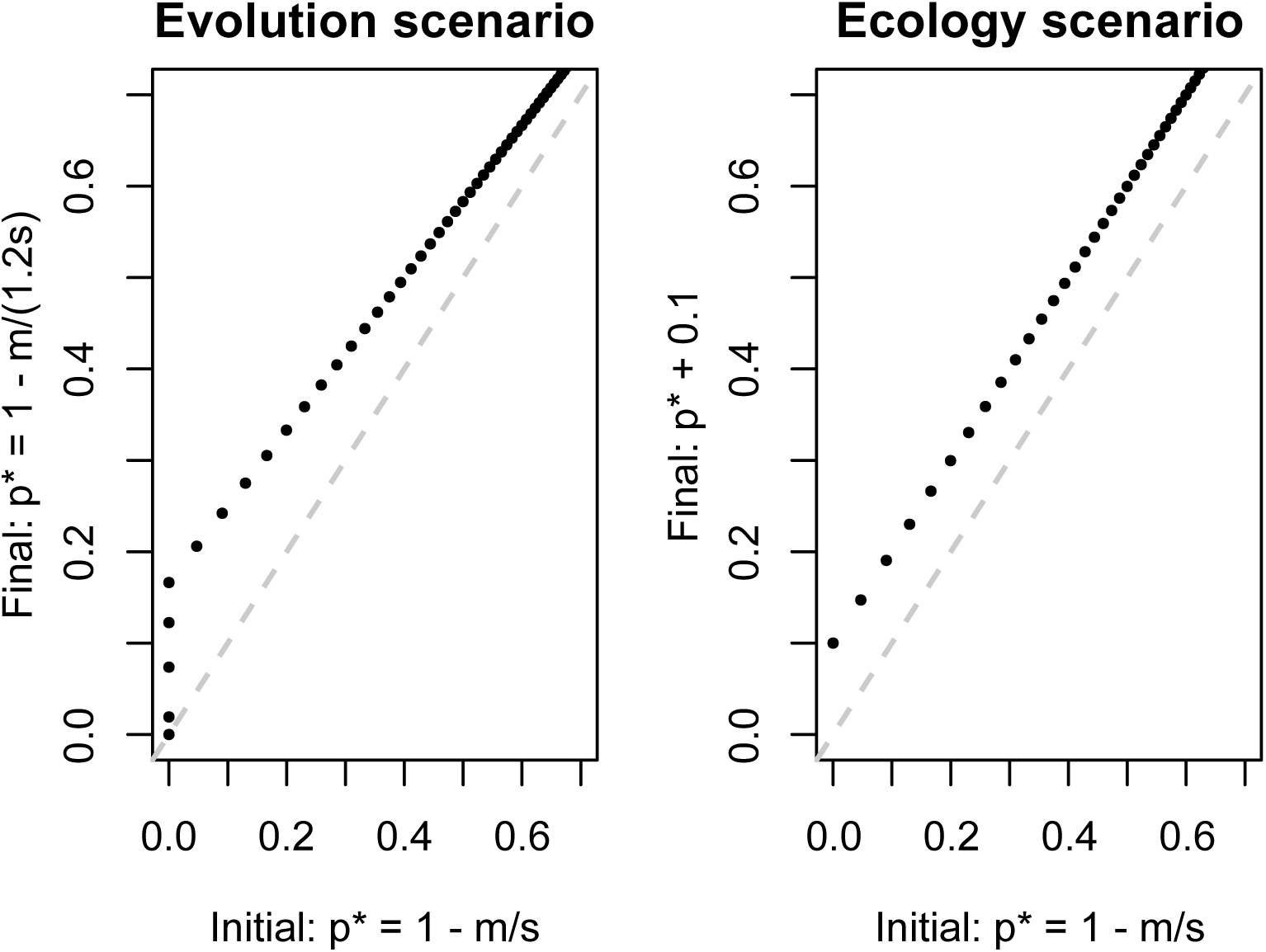
Simulation of two alternate scenarios for increasing human-specialist ancestry between two timepoints (initial, final) where p is human-specialist allele frequency, m is gene flow from forest populations, and s is the selection coefficient. Each dot represents some iteration across 2,000 values of *s* (0.001–1) with *m* constant at 0.01.

**Supplemental Figure 10.**
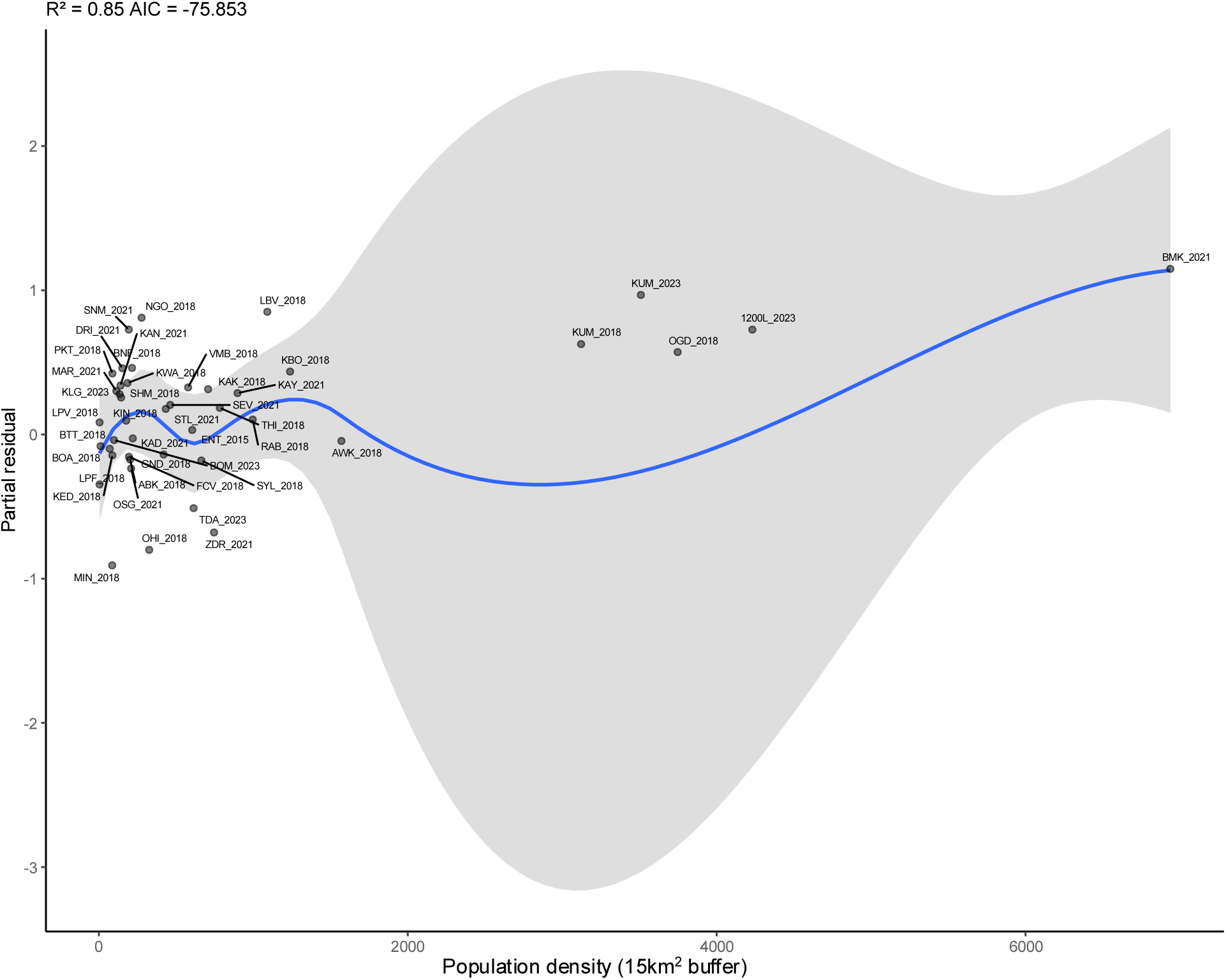
Partial residual plot from a GAM examining the relationship between population density (15 km buffer) and logit scaled human host preference, after accounting for smooth terms climate variables and batch included as a covariate. Blue line shows LOESS fit with grey 95% confidence interval.

**Supplementary Figure 11.**
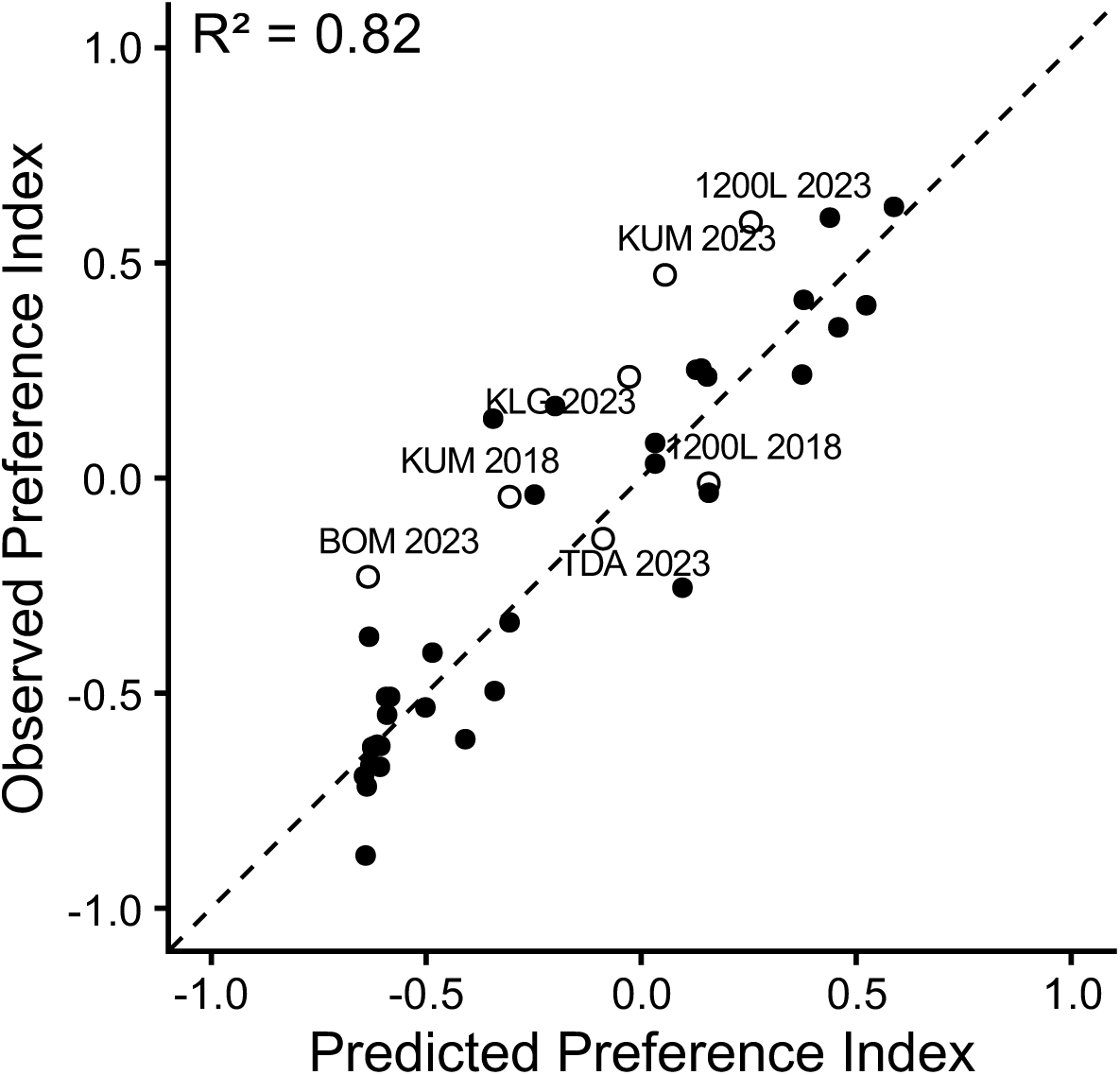
Correlation between the observed preference index and the predicted preference index from fastNGSadmix data at host-preference outlier regions. Dotted line shows 1:1 relationship. Open circles with labels show data from the current project, closed circles: Fifer et al. (2026) data.

**Supplementary Figure 12.**
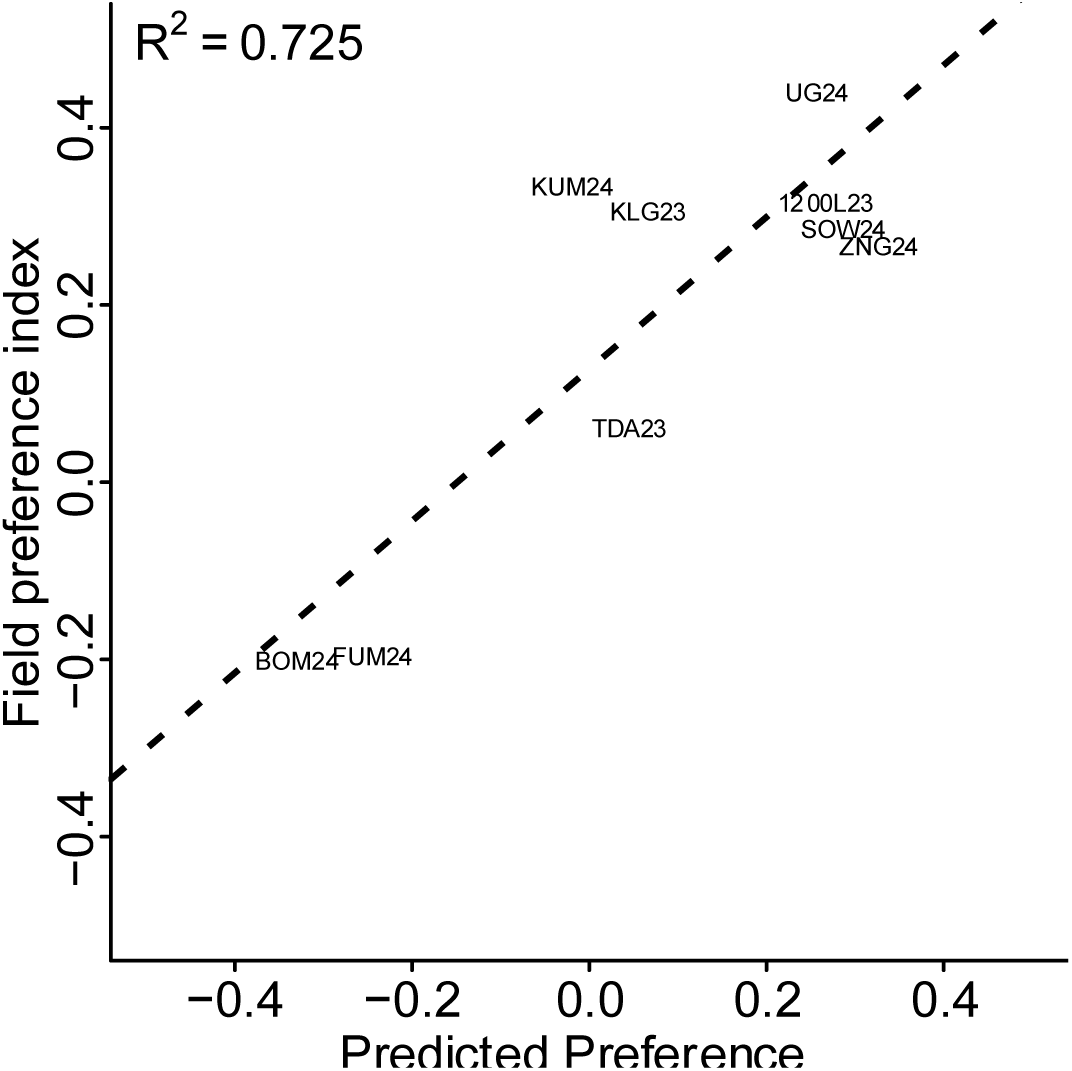
Correlation between the field preference index measured by paired animal-human baited BG traps and predicted preference index using fastNGSadmixture estimates at host-preference outlier regions. Dotted line shows 1:1 relationship.

**Supplementary Figure 13.**
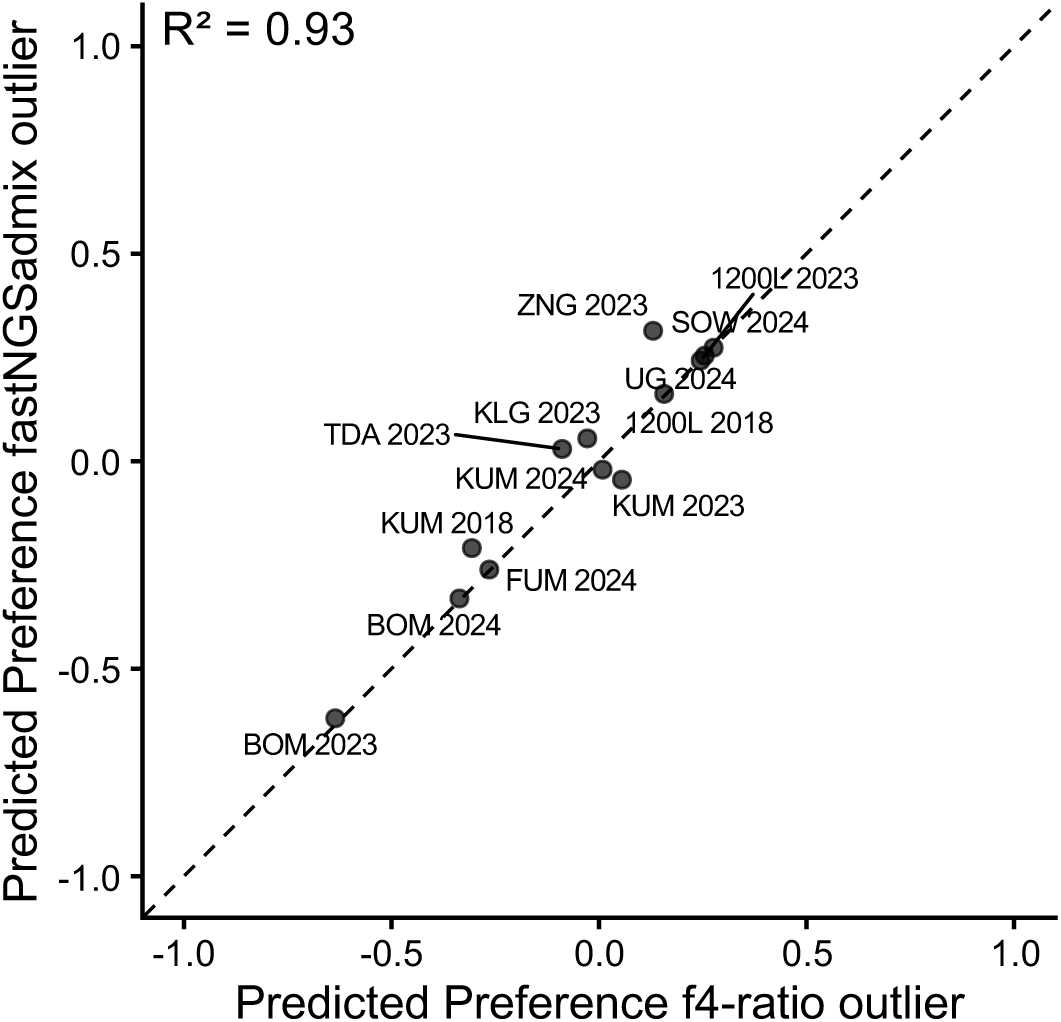
Correlation between the predicted preference index using either f4-ratio or fastNGSadmixture estimates at host-preference outlier regions. Dotted line shows 1:1 relationship.

**Supplementary Figure 14.**
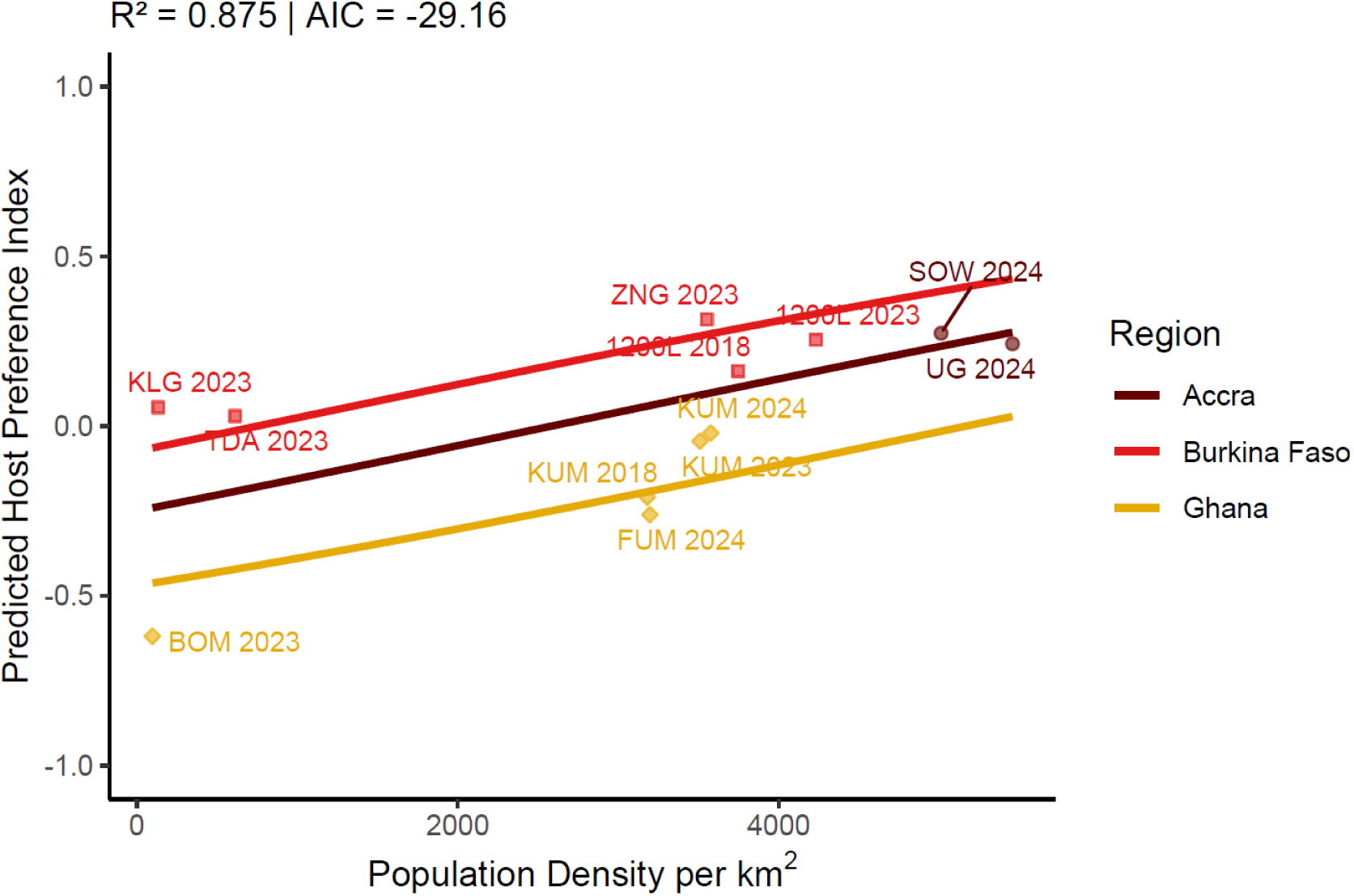
Relationship between human population density and predicted host preference index estimated using beta regression with geographic region included as a covariate. Fitted regression lines shown per region.

**Supplementary Figure 15.**
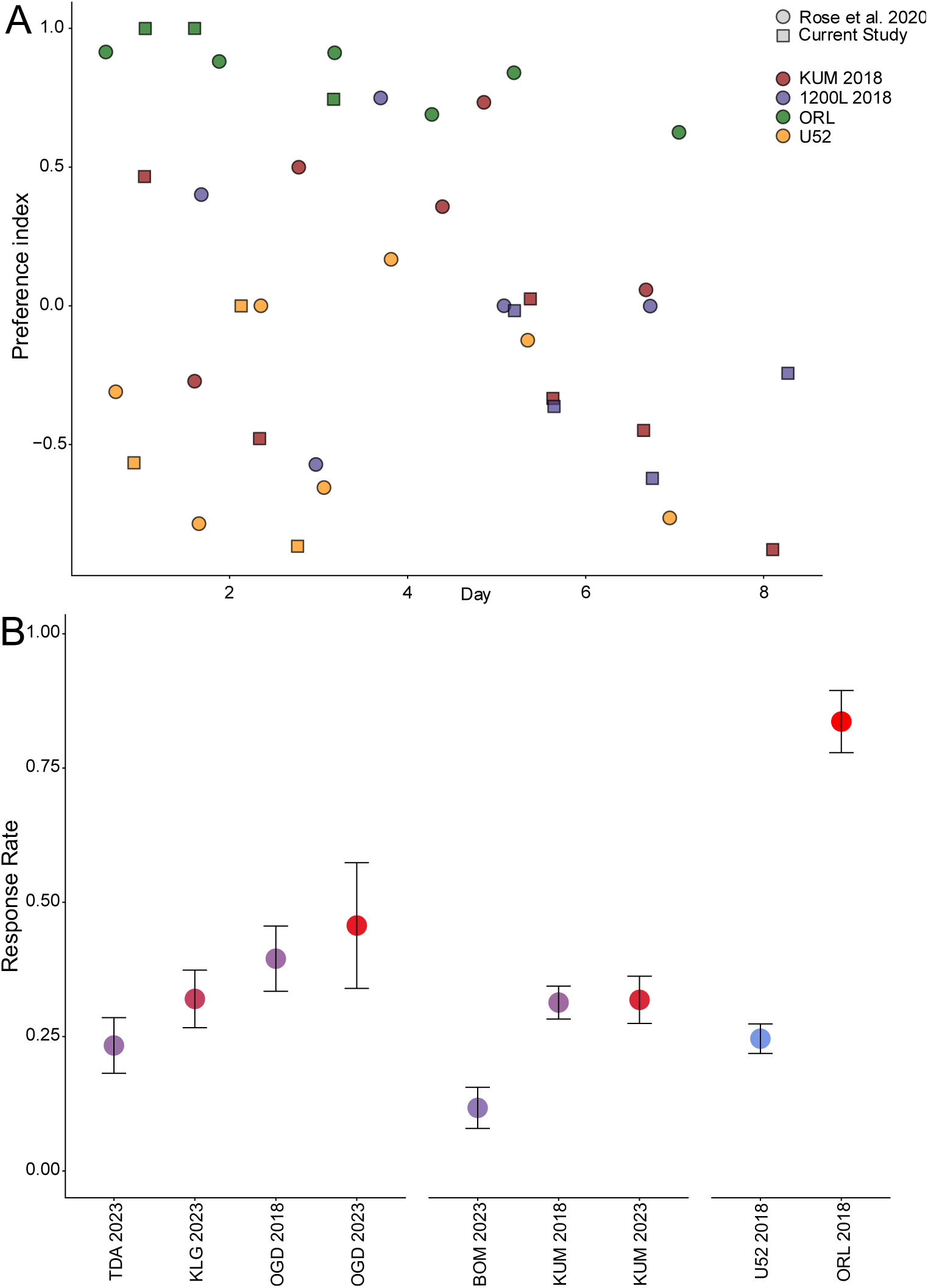
A) Comparison of the same colonies’ preference index between measurements taken with the olfactometer setup from Rose et al. (2020) versus the current study’s olfactometer. B) Mean response rate for all colonies from the current study with error bars showing 95% confidence interval.

**Supplementary Figure 16.**
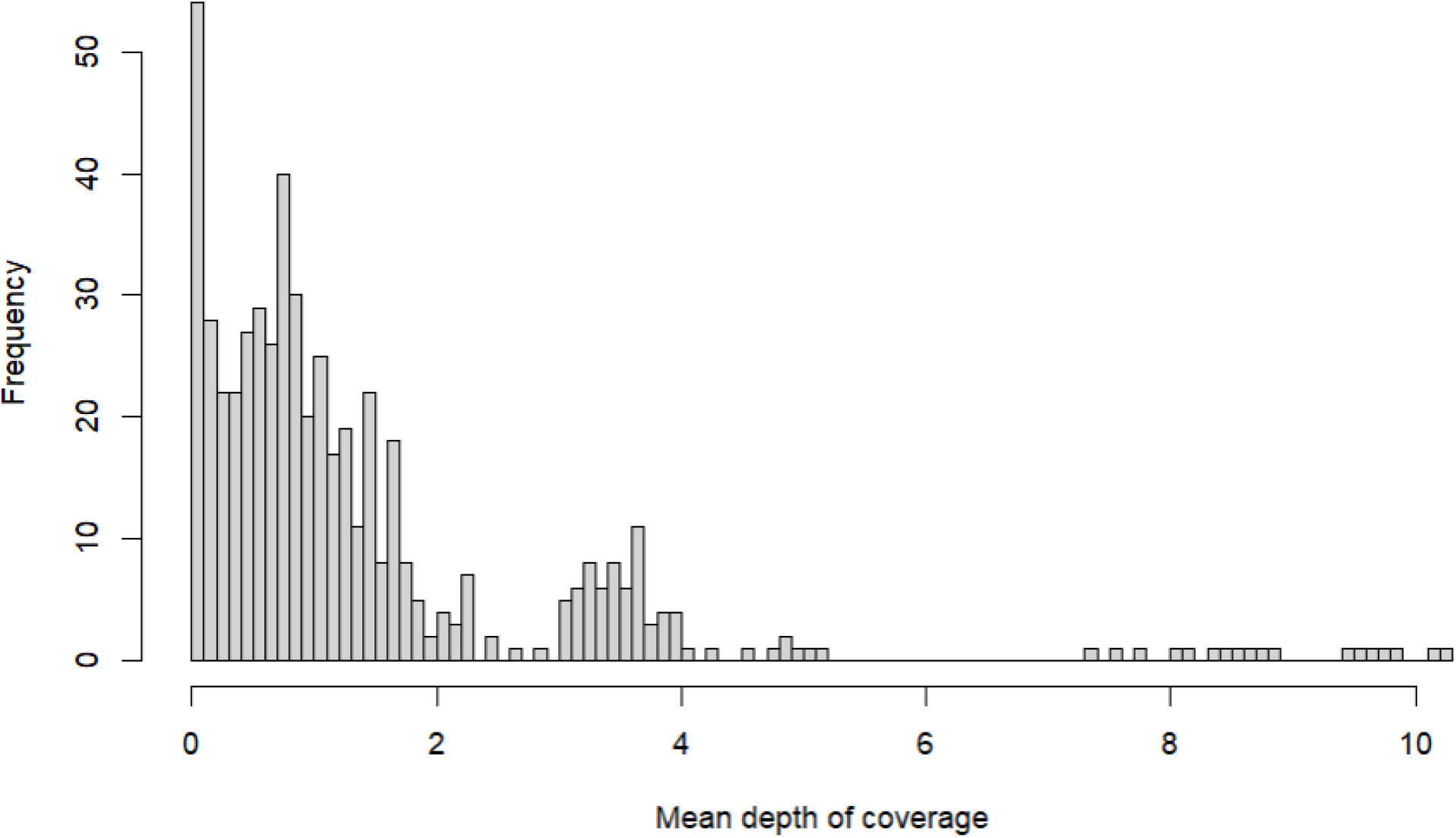
Distribution of mean chromosomal depth of coverage for all WGS data.

**Supplementary Figure 17.**
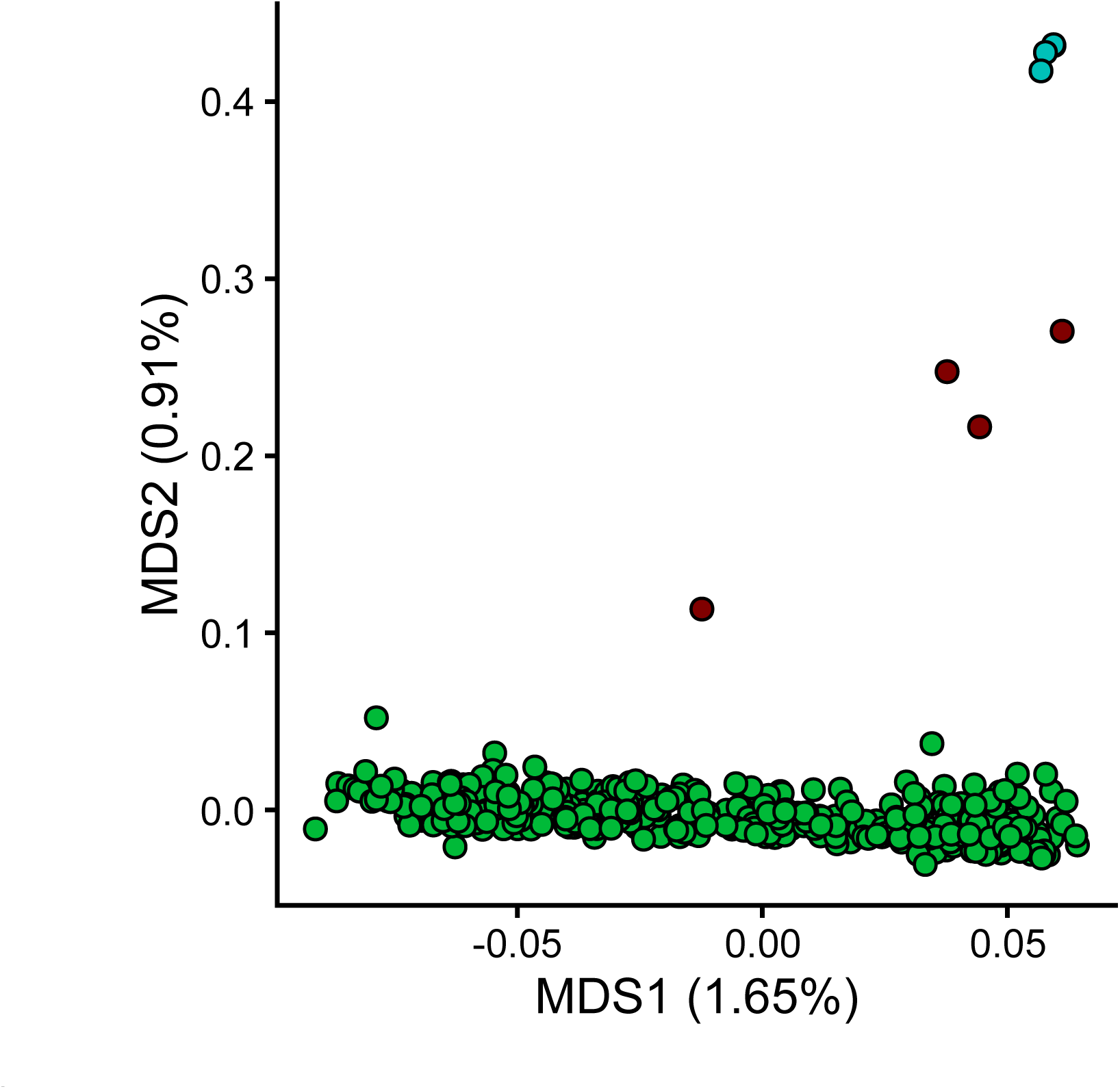
Principal Coordinates Analysis (PCoA) plot using a 1 -correlation transformation on the genetic covariance matrix of all Ghana and Burkina Faso samples with *Aedes mascarensis* (blue) included as outgroup to differentiate *Aedes aegypti* (green) from putatively non *Ae. aegypti* samples removed from analysis (red).

